# How repeated outbreaks drive the evolution of bacteriophage communication: Insights from a mathematical model

**DOI:** 10.1101/2020.04.29.068247

**Authors:** Hilje M. Doekes, Glenn A. Mulder, Rutger Hermsen

## Abstract

Communication based on small signalling molecules is widespread among bacteria. Recently, such communication was also described in bacteriophages. Upon infection of a host cell, temperate phages of the *Bacillus subtilis*-infecting SPbeta group induce the secretion of a phage-encoded signalling peptide, which is used to inform the lysis-lysogeny decision in subsequent infections: the phages produce new virions and lyse their host cell when the signal concentration is low, but favour a latent infection strategy, lysogenising the host cell, when the signal concentration is high. Here, we present a mathematical model to study the ecological and evolutionary dynamics of such viral communication. We show that a communication strategy in which phages use the lytic cycle early in an outbreak (when susceptible host cells are abundant) but switch to the lysogenic cycle later (when susceptible cells become scarce) is favoured over a bet-hedging strategy in which cells are lysogenised with constant probability. However, such phage communication can evolve only if phage-bacteria populations are regularly perturbed away from their equilibrium state, so that acute outbreaks of phage infections in pools of susceptible cells continue to occur. Our model then predicts the selection of phages that switch infection strategy when half of the available susceptible cells have been infected.

## Introduction

For several decades now, it has been recognised that communication between individuals is not limited to multi-cellular organisms, but is also common among microbes. The best-known example of microbial communication is bacterial *quorum sensing*, a process in which bacteria secrete signalling molecules to infer the local cell density and consequently coordinate the expression of certain genes [1, 2]. A wide variety of bacterial behaviours have been found to be under quorum-sensing control [2, 3], including bioluminescence [1], virulence [4], cooperative public good production [5, 6], and antimicrobial toxin production [7, 8]. Remarkably, it has recently been discovered that even some bacterial viruses (bacteriophages or phages for short) use signalling molecules to communicate [9]. Here, we use a mathematical model to explore the dynamics of this viral small-molecule communication system. We study under what conditions communication between phages evolves, and predict which communication strategies are then selected.

Bacteriophages of the *Bacillus*-infecting SPbeta group encode a small signalling peptide, named “arbitrium”, which is secreted when the phages infect bacteria [9]. These phages are *temperate* viruses, meaning that each time a phage infects a bacterium, it makes a life-cycle decision: to enter either (i) the *lytic* cycle, inducing an active infection in which tens to thousands of new phage particles are produced and released through host-cell lysis, or (ii) the *lysogenic* cycle, inducing a latent infection in which the phage DNA is integrated in the host cell’s genome (or episomally maintained) and the phage remains dormant until it is reactivated. This lysis-lysogeny decision is informed by the arbitrium produced in nearby previous infections: extracellular arbitrium is taken up by cells and inhibits the phage’s lysogeny-inhibition factors, thus increasing the propenstiy towards lysogeny of subsequent infections [9]. Hence, peptide communication is used to promote lysogeny when many infections have occurred. Similar arbitrium-like systems have now been found in a range of different phages [10]. Notably, these phages each use a slightly different signalling peptide, and do not seem to respond to the signals of other phages [9, 10].

The discovery of phage-encoded signalling peptides raises the question of how this viral communication system evolved. While the arbitrium system has not yet been studied theoretically, previous work has considered the evolution of lysogeny and of other phage-phage interactions. Early modelling work found that lysogeny can evolve as a survival mechanism for phages to overcome periods in which the density of susceptible cells is too low to sustain a lytic infection [11, 12]. In line with these model predictions, a combination of modelling and experimental work showed that selection pressures on phage virulence change over the course of an epidemic, favouring a virulent phage strain early on, when the density of susceptible cells is high, but a less virulent (*i*.*e*., lysogenic) phage strain later in the epidemic, when susceptible cells have become scarce [13, 14]. Other modelling work has shown that if phages, lysogenised cells, and susceptible cells coexist for long periods of time, the susceptible cell density becomes low because of phage exploitation, and less and less virulent phages are selected [15, 16].

Erez *et al*. [9] propose that the arbitrium system may have evolved to allow phages to cope with the changing environment during an epidemic, allowing the phages to exploit available susceptible bacteria through the lytic cycle when few infections have so far taken place and hence the concentration of arbitrium is low, while entering the lysogenic cycle when many infections have taken place and the arbitrium concentration has hence increased. This explanation resembles results for other forms of phage-phage interaction previously found in *Escherichia coli* -infecting phages [17, 18]. In the obligately lytic T-even phages, both the length of the latent period of an infection and the subsequent burst size increase if additional phages adsorb to the cell while it is infected – a process called *lysis inhibition* [18–20]. *In the temperate phage λ*, the the propensity towards lysogeny increases with the number of co-infecting virions, called the multiplicity of infection (MOI) [21]. In both cases, modelling work has shown that the effect of the number of phage adsorptions on an infection can be selected as a phage adaptation to host-cell density, as it allows phages to switch from a virulent infection strategy (*i*.*e*., a short latent period or a low lysogeny propensity) when the phage:host-cell ratio is low to a less virulent strategy (*i*.*e*., a longer latent period or higher lysogeny propensity) when the phage:host-cell ratio is high [22–24].

Here, we present a mathematical model to test if similar arguments can explain the evolution of small-molecule communication between viruses, and to explore the ecological and evolutionary dynamics of temperate phage populations that use such communication systems. We show that arbitrium communication can indeed evolve and that communicating phages consistently outcompete phages with non-communicating bet-hedging strategies. We however find that communication evolves under certain conditions only, namely if the phages regularly cause new outbreaks in substantial pools of susceptible host cells. Moreover, when communication evolves under such conditions, we predict that a communication strategy is selected in which phages use arbitrium to switch from a fully lytic to a fully lysogenic strategy when approximately half of all susceptible cells have been infected.

## Methods

### Model

Extending earlier models (*e*.*g*., [11, 13, 16, 24]), we use ordinary differential equations to describe a well-mixed system consisting of susceptible bacteria, lysogenically infected bacteria (also called lysogens), free phages, and an arbitrium-like signalling peptide (Fig 1A). For simplicity, we consider phages that do not affect the growth of lysogenised host cells; susceptible bacteria and lysogens hence both grow logistically with the same growth rate *r* and carrying capacity *K*. Lysogens are spontaneously induced at a low rate *α*, after which they lyse and release a burst of *B* free phages per lysing cell. Free phage particles decay at a rate *δ* and adsorb to bacteria at a rate *a*. Adsorptions to lysogens result in the decay of the infecting phage, thus describing the well-known effect of superinfection immunity [25–29], while adsorptions to susceptible bacteria result in infections with success probability *b*. We consider the lytic cycle to be fast compared to both bacterial growth and the lysogenic cycle (*c*.*f*., [11, 13, 16, 24]), so that a lytic infection can be modelled as immediate lysis releasing a burst of *B* free phages. Since the genes encoding arbitrium production are among the first genes to be expressed when a phage infects a host cell [9, 10], each infection leads to an immediate increase of the arbitrium concentration *A* by an increment *c*. The lysis-lysogeny decision is effected by the current arbitrium concentration: a fraction *φ*(*A*) of the infections results in the production of a lysogen, while the remaining fraction (1 *− φ*(*A*)) results in a lytic infection. Arbitrium does not decay spontaneously in the model (since it is a small peptide, spontaneous extracellular degradation is considered to be negligible), but it is taken up by bacteria at a rate *u* (*e*.*g*., through general bacterial peptide importers such as OPP [9]), and then degraded intracellularly.

**Fig 1.**
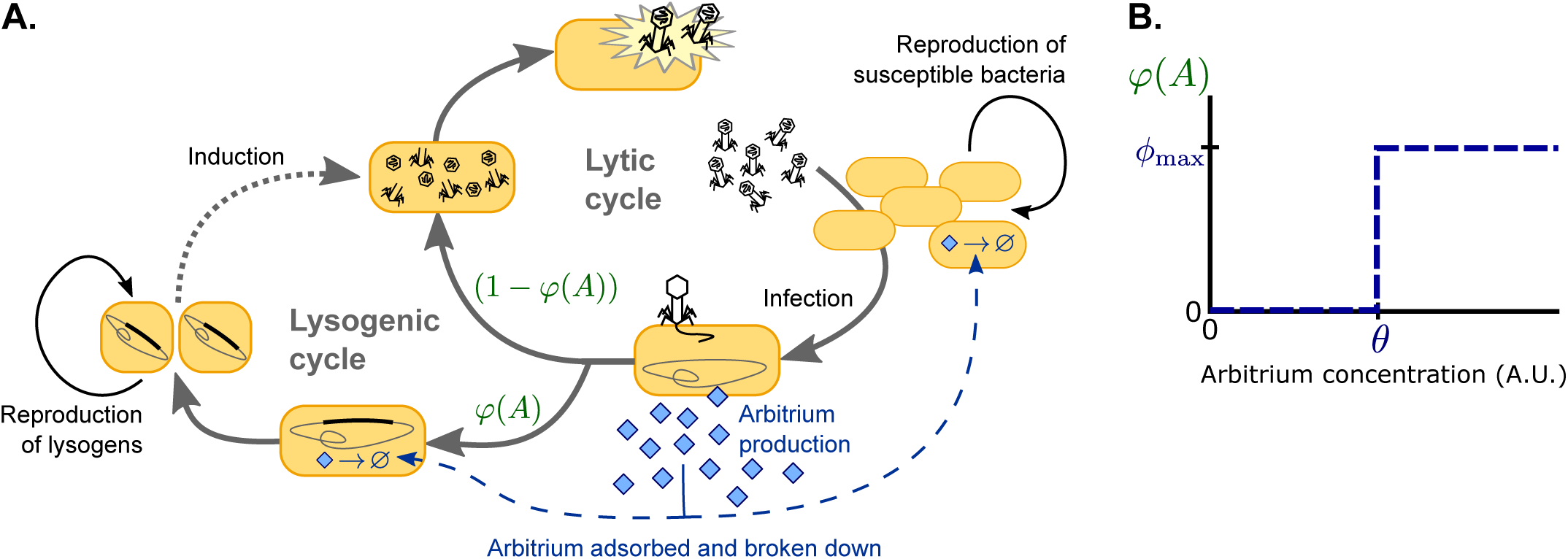
Model overview. A. Free phages infect susceptible bacteria, at which point a fixed amount of arbitrium is produced. This arbitrium is taken up and degraded by susceptible cells and lysogens. Upon infection, a cell enters the lysogenic cycle with propensity *φ*(*A*), or the lytic cycle with propensity (1 *− φ*(*A*)); the lysogeny propensity *φ*(*A*) depends on the current arbitrium concentration. The lytic cycle leads to immediate lysis of the host cell and release of a burst of new virions. In the lysogenic cycle, the phage remains dormant in the lysogen population, which grows logistically with the same rate as the susceptible cell population. Lysogens are spontaneously induced at a low rate, at which point they re-enter the lytic cycle. B. In communicating phages, the lysogeny propensity *φ*(*A*) is modelled by a step-function characterised by two phage characteristics: *θ*, the arbitrium concentration above which the phage increases its lysogeny propensity, and *ϕ*_max_, the lysogeny propensity of the phage at high arbitrium concentration.

Consider competing phage variants *i* that differ in their (arbitrium-dependent) lysogeny propensity *φ*_*i*_(*A*). The population densities of susceptible bacteria *S*, phage particles *P*_*i*_ and corresponding lysogens *L*_*i*_, and the concentration of arbitrium *A* can then be described by:

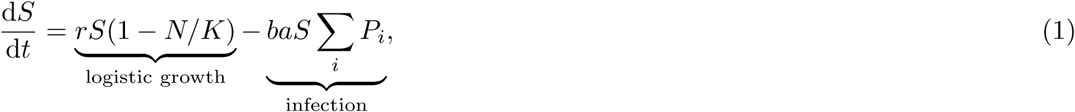

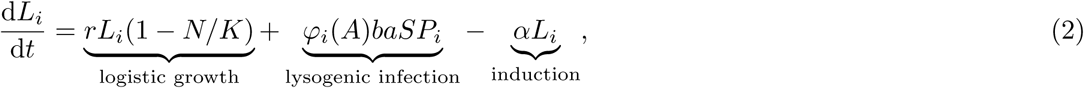

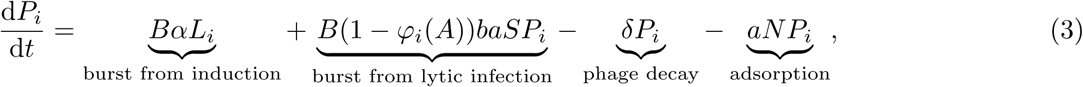

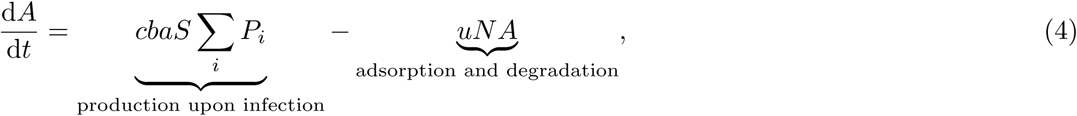

where *N* = *S* +Σ_*i*_ *L*_*i*_ is the total density of bacteria.

We study two scenarios for the lysis-lysogeny decision: (i) the arbitrium concentration does not affect the lysis-lysogeny decision; each phage variant has a constant lysogeny propensity *ϕ*_*i*_, and (ii) the arbitrium concentration does affect the lysis-lysogeny decision; each phage variant causes lytic infection when the arbitrium concentration is low, but switches to some lysogeny propensity *ϕ*_max__*i*_ when the arbitrium concentration exceeds the phage’s response threshold *θ*_*i*_ (Fig 1B). Though omitted from Eq 1–4 for readability, mutations between phage variants were included in the model (Supplementary Text S1.1). Under scenario (ii), mutations in *ϕ*_max_ and *θ* are implemented as independent processes.

### Parameters

In total, the model (Eq 1–4) has 9 parameters. As far as we are aware, none of these have been estimated for phages of the SPBeta group, but many have been measured for other phages, most of which infect *E*. *coli* [13, 30–34]. We first nondimensionalised the equations, which reduced the number of parameters to 5, and then derived estimates for these parameters using the data from other phages (Supplementary Text S1.3-4). To account for the uncertainty in these estimates, we analysed the model for a broad range of parameter values (Supplementary Table S1) to confirm that the results hold in general.

Numerical integration and analysis were performed in Matlab R2017b, using the built-in ODE-solver ode45. Scripts are available from the corresponding author upon request.

## Results

### Evolution of the lysis-lysogeny decision and arbitrium communication requires perturbations away from equilibrium

A common approach to analysing ODE-models such as Eq 1–4 is to characterise the model’s equilibrium states *(c*.*f*., [11, 16, 35]). Such an analysis is provided in Supplementary Text S2. However, we will here argue that to understand the evolution of arbitrium communication, and the lysis-lysogeny decision in general, considering the equilibrium states is insufficient.

Firstly, the function of the arbitrium system is to allow phages to respond to changes in the density of susceptible cells and phages as reflected in the arbitrium concentration. But when the population approaches an equilibrium state, the densities of susceptible cells and phages become constant, and so does the arbitrium concentration. Equilibrium conditions hence defeat the purpose of small-molecule communication such as the arbitrium system. Evolution of small-molecule communication must be driven by dynamical ecological processes, and hence can only be studied in populations that are regularly perturbed away from their ecological steady state.

Secondly, under equilibrium conditions natural selection can act on the lysis-lysogeny decision only if infections still take place, and hence lysis-lysogeny decisions are still taken. We argue that this is unlikely. If the phage population is viable (*i*.*e*., if the parameter values are such that the phages proliferate when introduced into a fully susceptible host population), the model converges to one of two qualitatively different equilibria, depending on parameter conditions (Supplementary Text S2): either (i) susceptible host cells, lysogens and free phages all coexist, or (ii) all susceptible host cells have been infected so that only lysogens and free phages remain. The evolution of a constant lysogeny propensity in a host-phage population with a stable equilibrium of type (i) was recently addressed by Wahl *et al*., who show that under these conditions selection always favours phage variants with high lysogeny propensity (*i*.*e*., *ϕ* = 1) [16]. However, only a narrow sliver of parameter conditions permits a stable equilibrium of type (i) [24, 35], and when we estimated reasonable parameter conditions based on a variety of well-studied phages, we found that they typically lead to a stable equilibrium of type (ii) (Supplementary Text S2, parameter estimates based on [13, 30–34]). This is because phage infections tend to be highly effective: their large burst size and consequent high infectivity cause temperate phages to completely deplete susceptible host cell populations, replacing them with lysogens that are immune to superinfection [36, 37]. In that case, after a short epidemic no more infections take place, and competition between different phage variants ceases (see Supplementary Fig S1 for example dynamics). Hence there is no long term selection on the lysis-lysogeny decision, and studying its evolution in this state is pointless.

We therefore consider a scenario in which the phage and cell populations are regularly perturbed away from equilibrium. To do so, we expose the system of Eq 1–4 to a serial-passaging regime (*c*.*f*., [38–42]). We initialise the model with a susceptible bacterial population at carrying capacity (*S* = *K*) and a small density of phages, numerically integrate Eq 1–4 for a time *T*, then transfer a fraction of the phage population to a new population of susceptible bacteria at carrying capacity, and repeat this cycle to bring about a long series of epidemics. The phage population is diluted when it is transferred but the relative frequency of the different phage variants is kept constant, thus ensuring that the phage variants that were highly prevalent in the phage population at the end of an episode remain at a high relative frequency at the start of the new episode.

### In the absence of arbitrium communication, bet-hedging phages are selected with low constant lysogeny propensity

To form a baseline expectation of the evolution of the lysis-lysogeny decision under the serial-passaging regime, we first considered a population of phage variants that do not engage in arbitrium communication, but do differ in their constant lysogeny propensity *ϕ*_*i*_. Under typical parameter conditions (Supplementary Text S1.3 and Supplementary Table 1), each passaging episode starts with an epidemic in which the susceptible cell population is depleted, followed by a period in which the bacterial population is made up of lysogens only (Fig 2A, dynamics shown for a passaging episode length *T* = 12 h). The composition of the phage and lysogen populations initially changes over subsequent passaging episodes (Fig 2A), but eventually an evolutionarily steady state is reached in which one phage variant dominates the phage population (*ϕ* = 0.04; Fig 2B), confirming that the lysis-lysogeny decision is indeed under selection.

**Fig 2.**
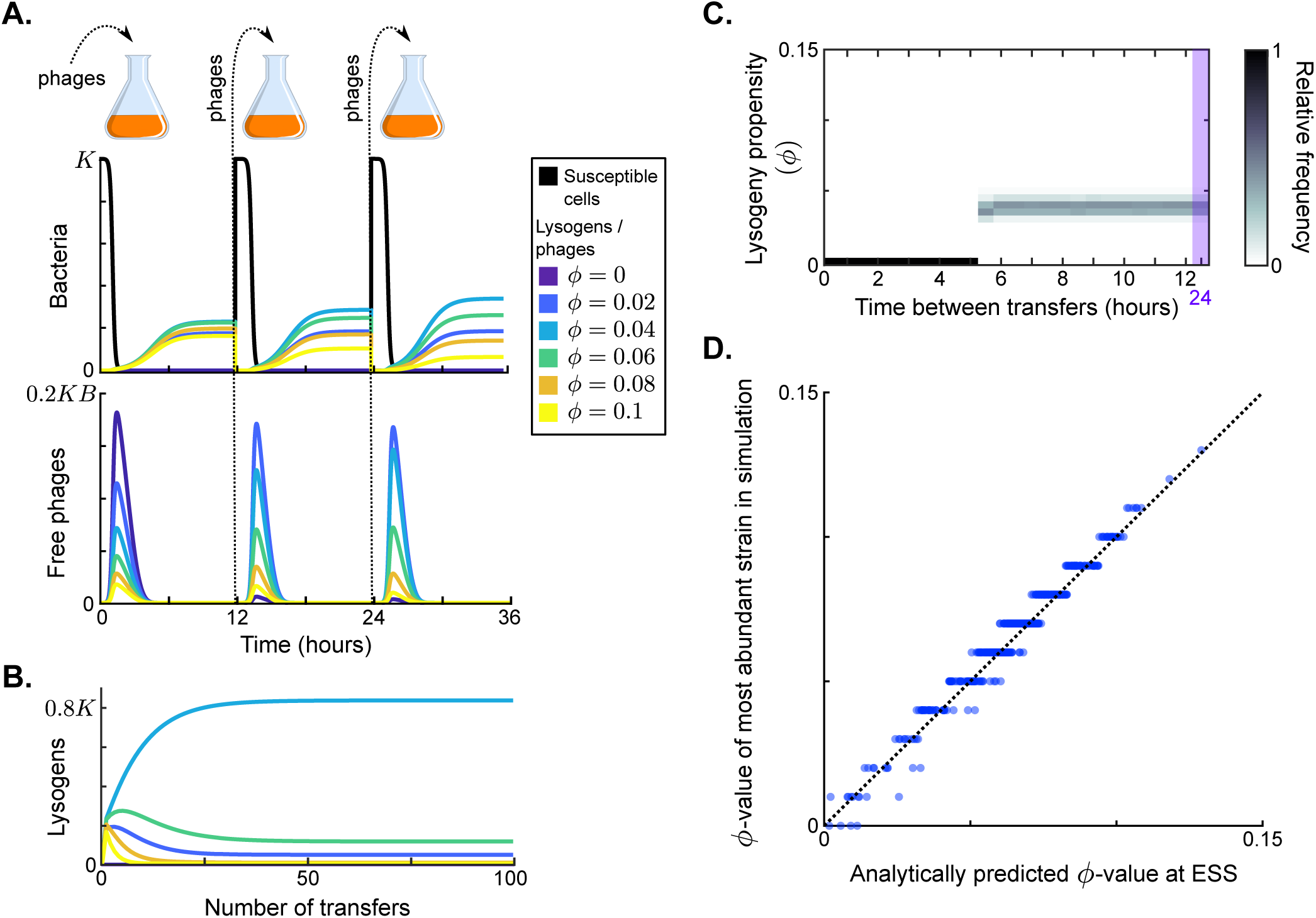
Results in the absence of phage communication. A. Short-term model dynamics under default parameter conditions (Supplementary Table S1) and a passaging episode duration of *T* = 12 hours. The model was initialised with a susceptible bacterial population at carrying capacity (*S* = *K*) and a low frequency of phages (Σ_*i*_ *P*_*i*_ = 10^*−*^5^^*KB*), and upon passaging the phages were diluted 100-fold. B. Long-term model dynamics for default parameter settings and *T* = 12 h. Over many passages, a single phage variant (*ϕ* = 0.04) is selected. C. Distribution of phage variants at evolutionarily steady state as a function of the time between passages, *T*. A total number of 101 phage variants was included, with lysogeny propensities varying between *ϕ*_1_ = 0 and *ϕ*_101_ = 0.5. When the interval between passages is short, the susceptible cells are not depleted during the rounds of infection and a fully lytic strategy (*ϕ* = 0) is selected. For larger values of *T*, however, a bet-hedging strategy with small but non-zero *ϕ*-value is selected (*ϕ ≈* 0.04). D. Parameter sweep results. The model was run with 500 sets of randomly sampled parameters, and for each run the most abundant *ϕ*-value in the population at evolutionarily steady state was plotted against the analytically predicted evolutionarily stable strategy (ESS; see Supplementary Text S3 and Box 1). The dotted line is the identity line. The analytically derived ESS is a good predictor of the simulation outcome.

The distribution of phage variants at evolutionarily steady state depends on the time between passages, *T* (Fig 2C). If this time is short (*T ≤* 5 h), the phage variant with *ϕ* = 0 dominates at evolutionarily steady state. This is an intuitive result: under these conditions phages are mostly exposed to environments with a high density of susceptible cells, in which a lytic strategy is favourable. Surprisingly, however, if the time between passages is sufficiently long (*T >* 5 h), the viral quasi-species at evolutionarily steady state always centres around the same phage variant, independent of *T* (*ϕ* = 0.04; Fig 2C). This result can be explained by considering the dynamics within a passaging episode (see Fig 2A and Supplementary Fig S1): Once the susceptible cell population has collapsed, free phages no longer cause new infections and are hence “dead ends”. New phage particles are then formed by reactivation of lysogens only, so that the distribution of variants among the free phages comes to reflect the relative variant frequencies in the lysogen population. Hence, when the time between passages is sufficiently long, the phage type that is most frequent in the sample that is eventually passaged is the one that is most frequent in the population of lysogens (Supplementary Fig S1). Under default parameter conditions, these are the phages with a low lysogeny propensity of *ϕ* = 0.04.

To examine how these results depend on the model parameters, we determined which phage variant was most abundant at evolutionarily steady state for 500 randomly chosen parameter sets (see Supplementary Table S1 for parameter ranges), always using a long time between passages (*T* = 24 h). The selected *ϕ*-values for all parameter settings lie between *ϕ* = 0 and *ϕ* = 0.12 (*y*-axis of Fig 2D). We can hence conclude that selection favours phages with low but usually non-zero lysogeny propensities. These phages employ a bet-hedging strategy: throughout the epidemic they “invest” a small part of their infection events in the production of lysogens, such that they are maximally represented in the eventual lysogen population.

To better understand how the lysogeny propensity *ϕ* that is selected depends on parameter values, we derived an analytical approximation for the evolutionarily stable strategy (ESS) under the serial-passaging regime (Supplementary Text S3.1-2). Because the phage dynamics during an epidemic affect the dynamics of the susceptible cells and *vice versa*, phage fitness is frequency dependent and the ESS is not found by a simple optimisation procedure, but by identifying the particular *ϕ*-value, denoted *ϕ*^*∗*^, that maximises phage fitness given that this strategy *ϕ*^*∗*^ itself shapes the dynamics of the epidemic (Box 1). We find that the ESS can be approximated by the surprisingly simple expression

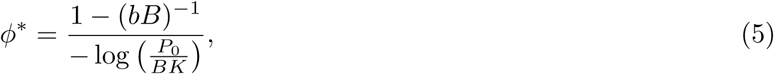

where *P*_0_ is the density of phages at the start of a passaging episode. Since *P*_0_*/*(*BK*) ≪ 1, the logarithm in the denominator of Eq 5 is negative. This approximation corresponds well with the results of the parameter sweep (Fig 2D), indicating that it indeed captures the most important factors shaping the evolution of the lysogeny propensity *ϕ*.

##### Box 1: Lysogeny propensity of the evolutionarily stable strategy (ESS)

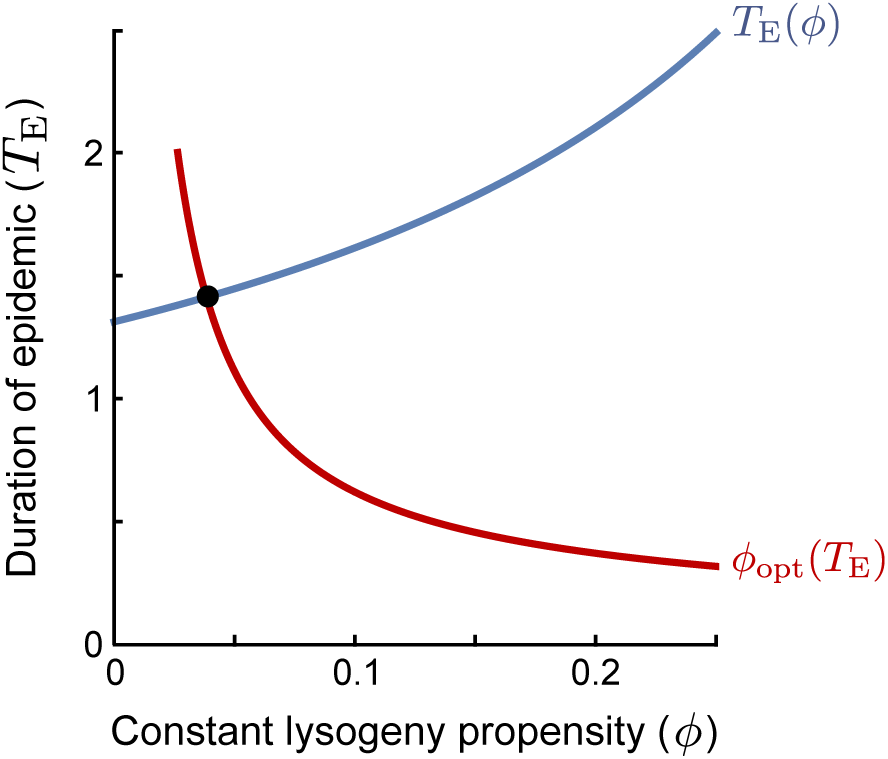

An evolutionarily stable strategy (ESS) is a strategy that cannot be invaded by any other strategy. In the context of the lysogeny propensity *ϕ*, it is the value *ϕ*^*∗*^ such that a population currently dominated by a phage with *ϕ* = *ϕ*^*∗*^ cannot be invaded2 by any phage variant with a different *ϕ*-value. A phage variant with *ϕ* = *ϕ*_*i*_ invading in a resident population with the same *ϕ* = *ϕ*_*i*_ always grows exactly like the resident. If this is the best possible invader, any other phage variant must perform worse than the resident and cannot invade. Hence, the ESS is the *optimal response to itself*. However, we still have to define what it means to be the “best possible invader” under the serial-passaging regime. Note that if the time between passages is sufficiently long, phages are selected on their ability to produce lysogens during the active epidemic (see Main Text). The optimal invader is hence the phage variant that, when introduced at a very low frequency, produces the most lysogens *per capita* between time *t* = 0 and the time that the susceptible cell population collapses, *T*_E_. The *ϕ*-value of the optimal invader depends on *T*_E_ (red line in plot): if the epidemic phase is short, lysogens have to be produced quickly and a high *ϕ*-value is optimal, while if the epidemic lasts longer, phages can profit more from lytic replication and a lower *ϕ*-value is optimal. In turn, however, the duration of the epidemic *T*_E_ depends on the lysogeny propensity *ϕ* of the resident phage population (blue line in plot): phages with a lower value of *ϕ* replicate more rapidly and hence cause an earlier collapse of the susceptible population. The ESS is the value *ϕ*^*∗*^ that is optimal given the collapse time *T*_E_(*ϕ*^*∗*^) that results when *ϕ*^*∗*^ itself is the resident strategy. Graphically, this value can be identified as the intersection of *T*_E_(*ϕ*) and *ϕ*_opt_(*T*_E_) (the red and blue lines).

Eq 5 shows that the ESS depends on the initial phage density in a passaging episode, *P*_0_, relative to the burst size *B* and maximal host-cell density *K*, and the effective burst size *bB*, which represents the expected number of progeny phages per phage that adsorbs to a susceptible bacterium. The ESS *ϕ*^*∗*^ decreases with the dilution factor of the phages upon passage (*i*.*e*., with lower *P*_0_). On the other hand, *ϕ*^*∗*^ increases with the effective burst size *bB* (note that (*bB*)^*−*^1^^ decreases when (*bB*) increases). Both effects can be intuitively understood by considering how these factors affect the duration of the epidemic, *T*_E_. If the phage density is low at the start of a passaging episode or if the phages have a small effective burst size, it takes a while before the phage population has grown sufficiently to cause the susceptible population to collapse. Since a lytic strategy is favoured early in the epidemic, when the susceptible cell density is still high, a longer epidemic favours phages with lower values of *ϕ* (see the red line in the figure in Box 1). On the other hand, if the initial phage density is high or if the phages have a high effective burst size, the susceptible cell population collapses quickly, phages have a much shorter window of opportunity for lysogen production and hence phages with higher *ϕ*-values are favoured.

### If arbitrium communication is included, communicating phages are selected that switch from a fully lytic to a fully lysogenic strategy

Next, we included the possibility of arbitrium communication and let phage variants be characterised by two properties: their arbitrium response threshold, *θ*_*i*_, and their lysogeny propensity when the arbitrium concentration exceeds their response threshold, *ϕ*_max__*i*_ (see Fig 1B). We then again considered the dynamics of our model under a serial-passaging regime.

In Fig 3A, example dynamics are shown for three competing phage variants, all with *ϕ*_max_ = 1 but with different response thresholds *θ*_*i*_. The arbitrium concentration increases over the course of the epidemic, and the phage variants switch from lytic infection to lysogen production at different times because of their different response thresholds.

**Fig 3.**
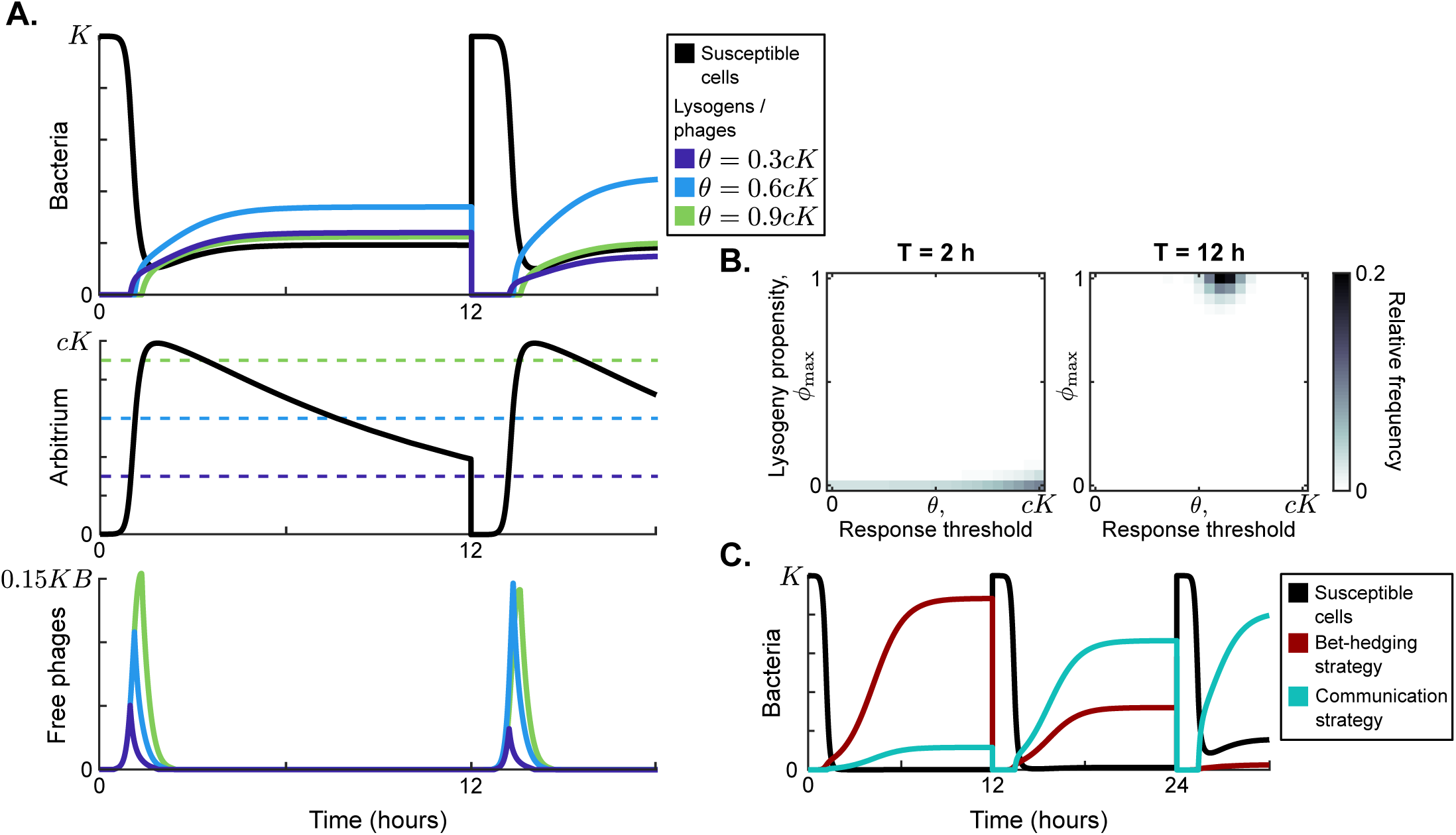
Model dynamics if phage communication is included in the model. A. Short-term dynamics for default parameter conditions (Supplementary Table S1) and the same serial-passaging regime as in Fig 2. This example shows the competition between three phage variants, all with *ϕ*_max_ = 1 but with varying response thresholds *θ*. B. Distribution of phage variants at evolutionary steady state for varying passaging episode durations *T*. In total 441 phage variants were included in this analysis, covering all combinations of *ϕ*_max_ between 0 and 1 and *θ* between 0 and *cK* with step sizes 0.05 and 0.05*cK*, respectively. When the interval between passages is very short, again a fully lytic strategy (*ϕ*_max_ = 0) is selected. For longer times between passages, however, we consistently see that a strategy with *ϕ*_max_ = 1 and *θ ≈* 0.65*cK* dominates the population. The results shown for *T* = 2 h are representative for values of *T ≤* 4 h, while the results shown for *T* = 12 h represent results obtained for *T ≥* 5 h (see Supplementary Fig S2). C. Rapid invasion by “optimally” communicating phages into a population of phages with the “optimal” bet-hedging strategy. The bet-hedging phages have *ϕ* = 0.04 (see Fig 2C), while the communicating phages are characterised by *ϕ*_max_ = 1 and *θ* = 0.66*cK* (see panel C). The communicating phage is initialised at a frequency of 1% of the bet-hedging phage.

Note that the maximum arbitrium concentration obtained during a passaging episode is approximately *A* = *cK* (Fig 3A). This is because during the epidemic the dynamics of the susceptible cell density are mostly determined by infection events and not so much by the (slower) bacterial growth. Since the arbitrium concentration increases by an increment *c* every time a susceptible cell is infected, the infection of all initial susceptible cells will result in an arbitrium concentration of *A* = *cK* (assuming that the degradation of arbitrium is also slow and can be ignored during the growth phase of the epidemic). The arbitrium concentration during the early epidemic then is a direct reflection of the fraction of susceptible cells that have so far been infected.

To study the evolution of arbitrium communication, we again considered the distribution of phage variants at evolutionary steady state for varying values of the time between passages, *T* (Supplementary Fig S2). Similar to the results shown in Fig 2C, we find two regimes (Fig 3B and Supplementary Fig S2): if the time between passages is short (*T <* 5 hours, illustrated by *T* = 2 h in Fig 3B)), selection favours phage variants that only cause lytic infections (*ϕ*_max_ = 0); if the time between passages is sufficiently long (*T ≥* 5 h, illustrated by *T* = 12 h in Fig 3B) the phage population is dominated by variants with *ϕ*_max_ = 1 and *θ ≈* 0.65*cK*. Hence, if the time between passages is sufficiently long, phage variants are selected that switch from a completely lytic to a completely lysogenic strategy when the arbitrium concentration exceeds a certain threshold.

In the simulations of Fig 3B, phage variants could have emerged that use the bet-hedging strategy found in the absence of communication (in phage variants with *θ* = 0, the lysogeny propensity is always *ϕ*_max_, independently of the arbitrium concentration), but this did not happen. This suggests that any bet-hedging phage variants were outcompeted by variants that do use arbitrium communication. To confirm this observation, we simulated a competition experiment between the bet-hedging phage variant that was selected in the absence of communication and the communicating variant selected when arbitrium dynamics were included (Fig 3C). The communicating phage quickly invades on a population of bet-hedging phages and takes over the population, confirming that communication is indeed favoured over bet-hedging.

### Evolved phages switch from the lytic to the lysogenic life-cycle when approximately half of the susceptible cells have been infected

To study how the evolution of phage communication depends on phage and bacterial characteristics, 500 simulations were performed with randomly sampled sets of parameter values (Supplementary Table S1), using a long time between serial passages (*T* = 24 h). For each simulation, we determined which phage variant was most prevalent at evolutionary steady state. Although we varied the parameter values over several orders of magnitude, the most prevalent phage variant had a lysogeny propensity of *ϕ*_max_ = 1 and a response threshold of *θ* = 0.5*cK* or *θ* = 0.6*cK* in almost all simulations (Fig 4A). Hence, over a broad range of parameter values, phages are selected that use the arbitrium system to switch from a fully lytic to a fully lysogenic strategy (*i*.*e*., *ϕ*_max_ = 1). This suggests that over the course of an epidemic, there is an initial phase during which the lytic strategy is a “better” choice (*i*.*e*., produces the most progeny on the long run), while later in the epidemic the production of lysogens is favoured and residual lytic infections that would results from a lysogeny propensity *φ <* 1 are selected against.

**Fig 4.**
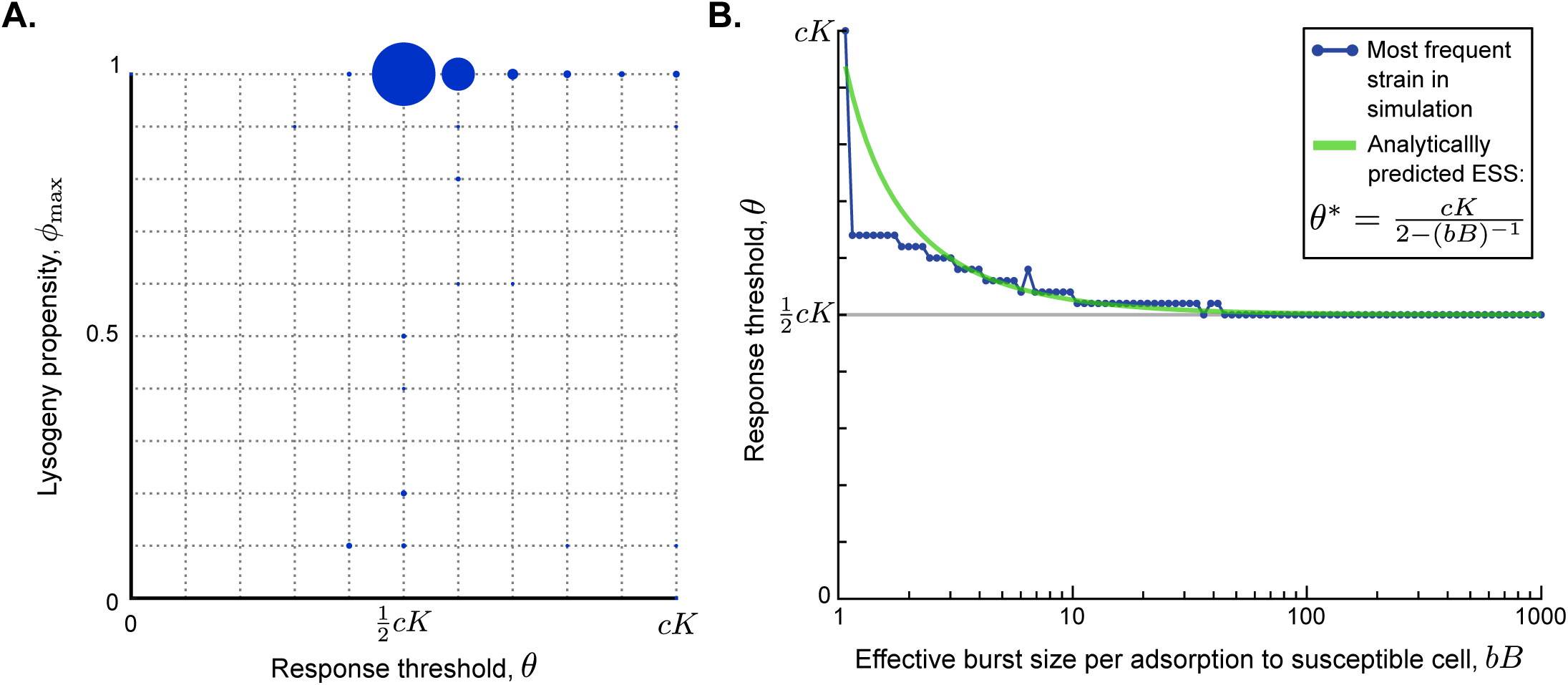
Parameter dependance of the selected values of *ϕ*_max_ and *θ*. A. Parameter sweep results. 500 simulations were run with randomly sampled parameters (Supplementary Table S1). In each simulation, 121 phage variants were included, covering all combinations of *ϕ*_max_ = 0 to *ϕ*_max_ = 1 and *θ* = 0 to *θ* = *cK* with step sizes 0.1 and 0.1*cK*, respectively. The size of the circles corresponds to the number of simulations that yielded that particular phage variant as most abundant at evolutionary steady state. B. Analytically predicted *θ*-value as a function of the effective burst size per adsorption to a susceptible cell, *bB*), and most abundant phage variant found in a simulation with varying *bB* but otherwise default parameter values, *T* = 24 h, *ϕ*_max_ = 1 and *θ* = 0, 0.02*cK, …, cK*. The range on the x-axis is equal to the range sampled in the parameter sweep. The analytically derived evolutionarily stable *θ*^*∗*^ is a good prediction for the response threshold selected in the simulations, especially for phages with high effective burst size.

To better understand the intriguing consistency in *θ*-values found in the parameter sweep, we used a similar approach as before to analytically derive an approximation for the response threshold *θ*^*∗*^ of the evolutionarily stable strategy (Supplementary Text S3.3). Again, we find a surprisingly simple expression for the ESS:

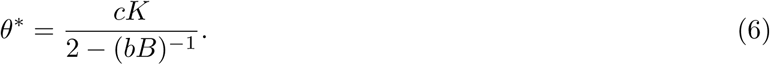

Note that the expression in Eq 6 again depends on the effective burst size *bB*, which is an indicator of the phage’s infectivity. The evolutionarily stable response threshold *θ*^*∗*^ declines as the effective burst size increases, converging to a value of 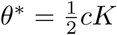 for highly infective phages (Fig 4B, green line). The same result was found for simulations of the competition between phage variants with different *θ*-values under different effective burst sizes (Fig 4B, blue dots). Eq 6 hence provides a good prediction for the response threshold value that is selected over evolutionary time, especially for phages with high effective burst size (Fig 4B).

The result in Eq 6 can be further understood biologically. Remember that the arbitrium concentration during the epidemic varies between *A* = 0 and *A* = *cK*, and is a reflection of the fraction of susceptible cells that have so far been infected. It makes sense that the evolutionarily stable response threshold causes phages to switch infection strategy somewhere in the middle of the epidemic: if a phage variant switches to the lysogenic strategy too early, its free phage population does not expand enough to compete with phages that switch later, but if it switches too late, the susceptible-cell density has decreased to such a degree that the phage has missed the window of opportunity for lysogen production. The ESS results from a balance between the fast production of phage progeny during the initial lytic cycles and the eventual production of sufficient lysogens. For phages with a high effective burst size, this balance occurs around the time that half of the available susceptible cells have been infected. Phages with lower effective burst size are however predicted to switch later, because these phages need to invest a larger portion of the available susceptible cells in the production of free phages to produce a sufficient pool of phages that can later form lysogens. Note, however, that the range of biologically reasonable effective burst sizes includes many high values (range of *x*-axis in Fig 4B, Supplementary Table S1), *i*.*e*., many real-life phages have high infectivity. Hence, for natural phages in general, we predict that if they evolve an arbitrium-like communication system, communication will be used to switch from causing mostly lytic to mostly lysogenic infections when in an outbreak approximately half of the pool of susceptible bacteria has been infected.

## Discussion

We have presented a mathematical model of a population of phages that use an arbitrium-like communication system, and used this model to explore the evolution of the lysis-lysogeny decision and arbitrium communication under a serial-passaging regime. When arbitrium communication was excluded from the model, we found that bet-hedging phages with relatively low lysogeny propensity were selected. But when arbitrium communication was allowed to evolve these bet-hedging phages were outcompeted by communicating phages. These communicating phages switch from a lytic strategy early in the epidemic to a fully lysogenic strategy when approximately half of the available susceptible cells have been infected.

The serial-passaging set-up of the model is crucial for the evolution of the lysis-lysogeny decision and arbitrium communication. This has two main reasons. Firstly, it ensures that the phages are regularly exposed to susceptible cells, thus maintaining selection pressure on the lysis-lysogeny decision. Because of their high infectivity (see Methods section and [31, 32]), most temperate phage outbreaks will completely deplete pools of susceptible bacteria, resulting in a bacterial population consisting of lysogens only in which the phage no longer replicates through infection [36, 37]. The bet-hedging strategy we found in the absence of phage communication is a mechanism to deal with these (self-inflicted) periods of low susceptible cell availability, consistent with earlier studies [12, 24]. Secondly, the serial-passaging set-up imposes a dynamic of repeated epidemics in which a small number of phages is introduced into a relatively large pool of susceptible cells. Such dynamics are necessary for the arbitrium system to function: the arbitrium concentration provides a reliable cue for a phage’s lysis-lysogeny decision only if it is low at the beginning of an epidemic and subsequently builds up to reflect the fraction of cells that have so far been infected.

Based on these considerations, we can stipulate which environments promote the evolution of small-molecule communication such as the arbitrium system. One major factor that can ensure a regular exposure to susceptible cells (the first requirement) is spatial structure. If phages mostly infect bacteria that are physically close to them, a global susceptible population can be maintained even though susceptible bacteria may be depleted in local environments [43]. Indeed, spatial structure has been shown to greatly influence phage evolution, for instance by promoting the selection of less virulent strains that deplete their local host populations more slowly [43–45]. For small-molecule communication to evolve, however, the phages would also have to undergo repeated, possibly localised, outbreak dynamics (the second requirement). Such dynamics could occur in structured meta-populations of isolated bacterial populations, between which the phages spread at a limited rate. Alternatively, phages might encounter large pools of newly susceptible bacteria if they escape superinfection immunity through mutation [46–48]. This scenario does however require the arbitrium concentration to be low when the mutation occurs, so that the signal again provides information about the number of susceptible cells available. This could happen either if the arbitrium produced during previous epidemics has been degraded once the mutation occurs, or if the mutation co-occurs with a second mutation that changes the phage’s signal specificity. This second argument might in part explain the large diversity of phage signalling peptides observed [9, 10].

The model presented in this paper allows us to put hypotheses about the arbitrium system to the test. For instance, it has been suggested that the arbitrium system would benefit from the production of arbitrium by lysogens, because phages thereby would be warned about the presence of neighbouring lysogens (which are immune to superinfection) [49]. Above we have argued, however, that under repeated epidemics, such as caused by serial passaging, selection on the lysis-lysogeny decision and arbitrium signalling is limited to the relatively short window of time in which all (locally) present susceptible cells become infected: afterwards no new infections occur and arbitrium therefore has no effect. During this short time window, the density of lysogens is still low, and any arbitrium produced by lysogens contributes little to the information already conveyed by arbitrium produced during infection events. Hence, our model predicts that, under repeated epidemics that completely deplete (local) pools of susceptible cells, the effects of arbitrium production by lysogens are likely very minimal.

Intriguingly, our model predicts that phages using small-molecule communication to coordinate their lysis-lysogeny decision would be selected to switch from a lytic to a lysogenic strategy once approximately half of the available susceptible bacteria have been lytically infected. This prediction warrants experimental testing. However, it also raises the question of how the phages would “know” at what bacterial density the susceptible population has been halved. For the arbitrium signal to carry information about the density of remaining susceptible cells, the initial concentration of susceptible bacteria has to be similar from outbreak to outbreak. While this was imposed in the serial-passaging regime used here, it is much less obvious that such a requirement would be met in natural environments. This limitation, however, is not unique to phage communication. For instance, it also complicates bacterial quorum sensing: as previously described, the concentration of a signalling molecule is often not determined by the local cell density only, but also depends on environmental properties [50, 51].

Next to the arbitrium system, several other examples of temperate phages affected by small signalling molecules have recently been described. For instance, the *Vibrio cholerae*-infecting phage VP882 “eavesdrops” on a quorum sensing signal produced by its host bacteria, favouring lytic over lysogenic infections when the host density is high [52], while in coliphages *λ* and T4 and several phages infecting *Enterococcus faecalis*, the induction of prophages, *i*.*e*., the lysogeny-lysis decision, is affected by bacterial quorum sensing signals [53–55]. Mathematical and computational modelling can help to better understand the ecology and evolution of these fascinating regulation mechanisms as well.

## Acknowledgments

We thank Rob J. de Boer for valuable discussions and comments on the manuscript. This work was supported by the Human Frontier Science Program, grant nr. RGY0072/2015 (http://www.hfsp.org/funding/research-grants).

## Author contributions

HMD conceived the study with input from RH. HMD and GAM performed and analysed the numerical simulations. HMD and RH derived the analytical results. RH supervised the study. HMD wrote the manuscript, with edits by GAM and RH.

## Supplementary Information

### S1 Model equations and Parameters

#### S1.1 Full model equations, including mutations

For readability, the model equations in the main text (Eq 1–4) did not include mutations between phage variants. Here, we present the full model equations, including mutations. We assume that mutations happen when new phage particles (and hence new copies of the phage’s genetic material) are formed prior to lysis of the host cell. Infection with a parent phage of variant *j* results in progeny phages of variant *i* with probability *µ*_*ij*_, where *µ*_*ii*_ is the probability that offspring of a parent phage of type-*i* has no mutations. The full model reads:

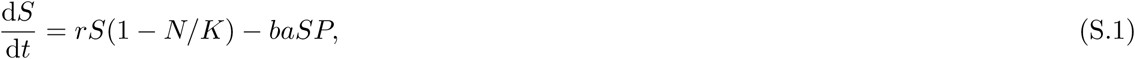

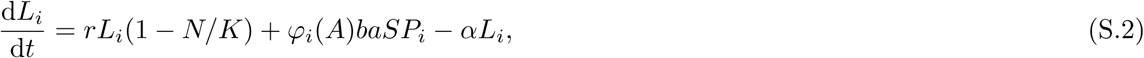

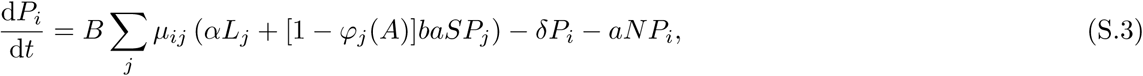

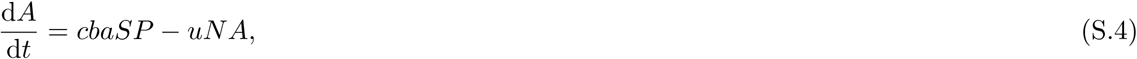

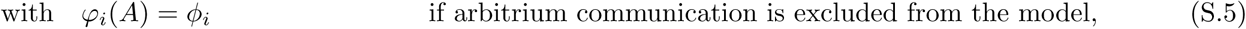

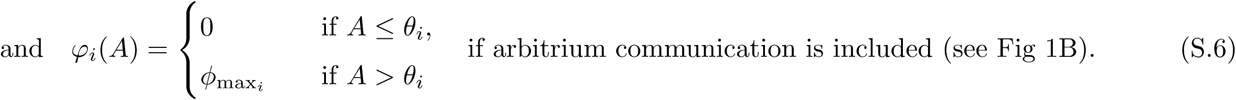

In the first part of the manuscript, where we exclude arbitrium communication from the model, phage variants are characterised by their constant lysogeny propensity *ϕ*_*i*_ (Eq S.5). Mutations between phage variants were implemented in a hill-climbing fashion, *i*.*e*., *µ*_*ij*_ = *µ >* 0 if *ϕ*_*j*_ is one step higher or lower than *ϕ*_*i*_, and *µ*_*ij*_ = 0 (*i I*= *j*) otherwise. Throughout this study, a value of *µ* = 5 *·* 10^−4^ was used. Varying *µ* affects the width of the quasispecies found, but not the actual outcome of the simulations.

In the second part of the manuscript, where we include the possibility of arbitrium communication, phage variants are characterised by two properties (Eq S.6): their arbitrium threshold *θ*_*i*_ and the lysogeny propensity obtained when the arbitrium concentration exceeds this threshold, *ϕ*_max__*i*_. Mutations in the values of *ϕ*_max_ and *θ* were implemented as independent processes, both happening in a hill-climbing fashion as explained above.

#### S1.2 Phage variants included in simulations

In the simulations of the restricted model (arbitrium communication excluded) shown in Fig 2C, a range of phage variants was included with *ϕ*_1_ = 0, *ϕ*_2_ = 0.005, *…, ϕ*_100_ = 0.495, *ϕ*_101_ = 0.5. In the simulations presented in Fig 2D, a range of phage variants was included of *ϕ*_1_ = 0, *ϕ*_2_ = 0.01, *…, ϕ*_100_ = 0.99, *ϕ*_101_ = 1.0.

In the simulations of the full model (arbitrium communication included) shown in Fig 3B and Supplementary Fig S2, 441 phage variants were included, representing all possible combinations of *ϕ*_max_ = 0, 0.05, *…*, 1 and *θ* = 0, 0.05*cK, …, cK*. In the simulations of Fig 4A, 121 phage variants were included representing all possible combinations of *ϕ*_max_ = 0, 0.1, *…*, 1 and *θ* = 0, 0.1*cK, …, cK*. In the simulations of Fig 4B, all phage variants had *ϕ*_max_ = 1, but they varied in *θ* = 0, 0.02*cK, …, cK*.

#### S1.3 Parameter reduction

In total, the model of Eq S.1-S.6 has 9 parameters (ignoring mutation probabilities). This number can, however, be reduced by non-dimensionalising the equations. We introduce the dimensionless variables

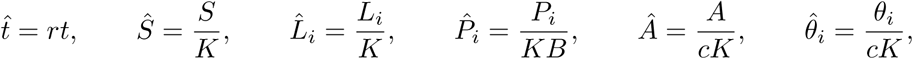

and define 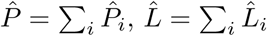, and 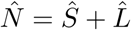. Let furthermore

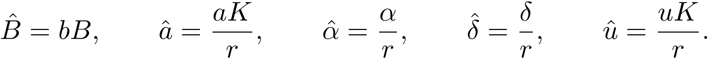

Using these new variables and parameters, the equations reduce to:

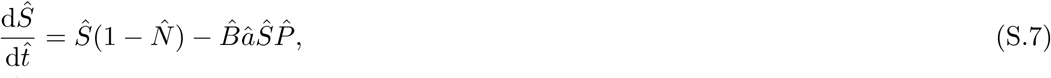

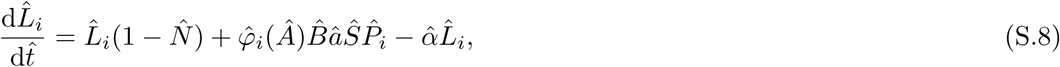

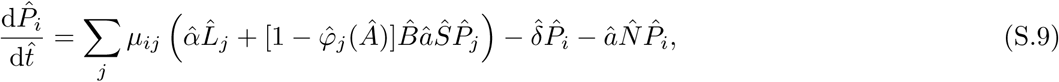

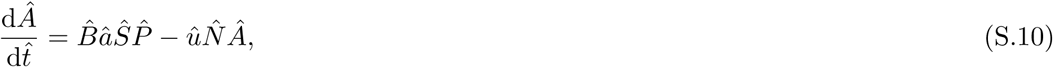

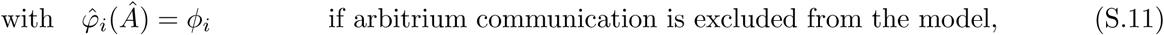

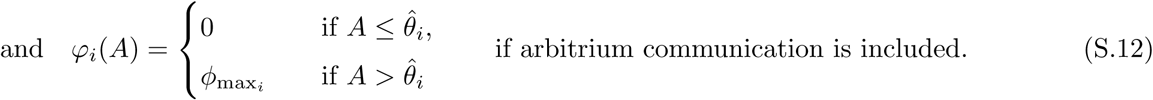

Five dimensionless parameters are left in Eq S.7–S.12: the effective burst size per adsorption of a phage to a susceptible cell, 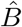, and the scaled rates *â*, 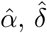, and 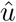. These non-dimensionalised equations are used throughout the rest of this supplementary text, unless stated otherwise, dropping the hats for convenience.

#### S1.4 Parameter values

As far as we are aware, no estimates exist for the parameters characterising phage phi3T. However, many parameters have been measured for a variety of phages infecting *Escherichia coli* [13, 30–34]. Based on these measurements, we derived reasonable ranges for the model parameters, and chose default parameter values (Supplementary Table 1). To study how our results depend on the model parameter values, parameter sweeps were performed, consisting of 500 simulations with parameter values randomly sampled from the broad parameter ranges (Supplementary Table 1). To ensure that low values of the parameters were well-represented, parameter values were sampled log-uniformly.

A small default value of the effective burst size was chosen to enhance the clarity of Fig 2A and Fig 3A,C; with a small effective burst size, the epidemic takes some time and competition between phage variants can be visualised more clearly. But much larger effective burst sizes were appropriately included in the parameter sweeps.

**Supplementary Table 1.**
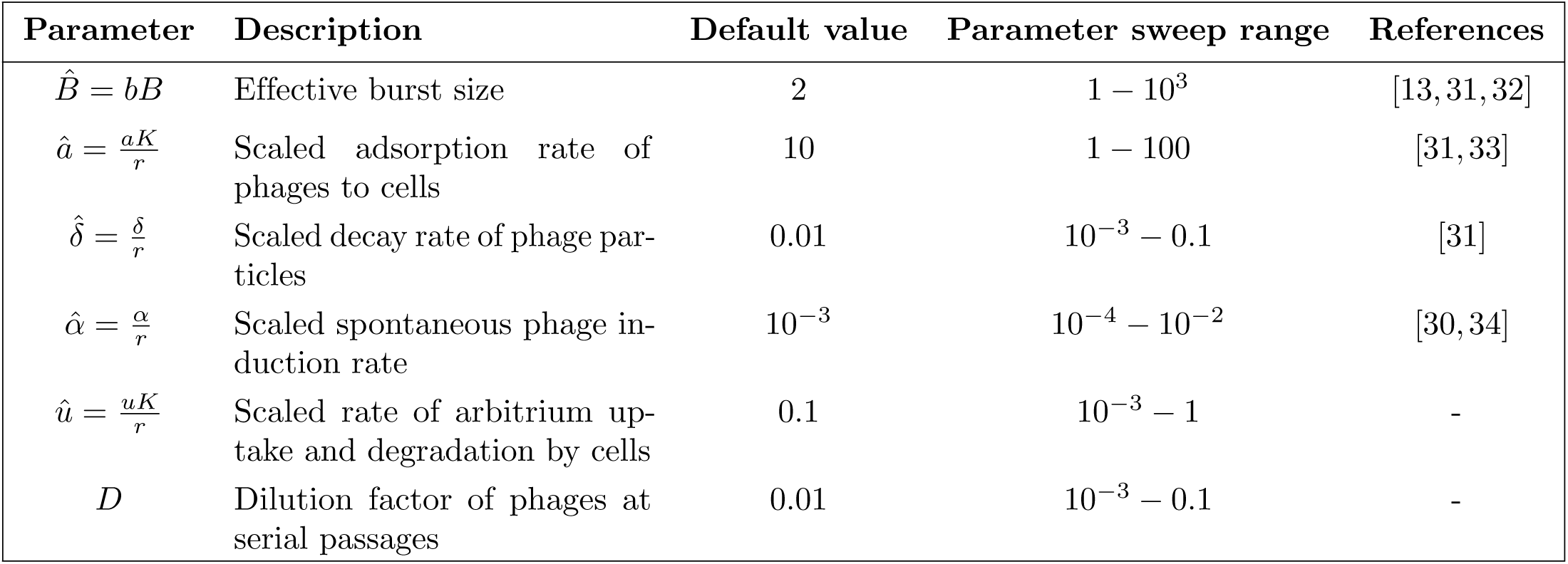
Model parameters.

| Parameter | Description | Default value | Parameter sweep range | References |
| --- | --- | --- | --- | --- |
| $\hat{B} = bB$ | Effective burst size | 2 | $1 - 10^3$ | [13, 31, 32] |
| $\hat{a} = \frac{aK}{r}$ | Scaled adsorption rate of phages to cells | 10 | $1 - 100$ | [31, 33] |
| $\hat{\delta} = \frac{\delta}{r}$ | Scaled decay rate of phage particles | 0.01 | $10^{-3} - 0.1$ | [31] |
| $\hat{\alpha} = \frac{\alpha}{r}$ | Scaled spontaneous phage induction rate | $10^{-3}$ | $10^{-4} - 10^{-2}$ | [30, 34] |
| $\hat{u} = \frac{uK}{r}$ | Scaled rate of arbitrium uptake and degradation by cells | 0.1 | $10^{-3} - 1$ | - |
| $D$ | Dilution factor of phages at serial passages | 0.01 | $10^{-3} - 0.1$ | - |

### S2 Equilibrium analysis

In this section, we find the dynamical equilibria existing in the model, and derive parameter conditions for their stability. This analysis provides us with a baseline expectation of the densities of phages, lysogens, and susceptible cells that the model converges to after sufficient time.

Equilibria of the model are found by equating Eq S.7–S.10 to zero and solving for the model variables. By the definition of equilibrium, the arbitrium concentration at equilibrium is constant:

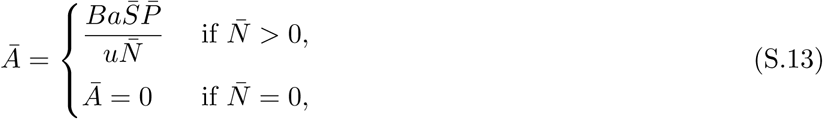

where the bar indicates equilibrium values. Because the equilibrium arbitrium concentration is constant, at equilibrium the different phage variants are characterised by their lysogeny propensity *φ*_*i*_(*Ā*) only, irrespective of whether arbitrium communication is included in the model or not. Hence, expressions for the model equilibria are the same in the absence and presence of arbitrium communication. Below, we derive these expressions, and determine under what conditions the different model equilibria are stable.

In the model, four qualitatively different types of equilibria can occur.

Firstly, there is a trivial equilibrium at 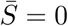, and 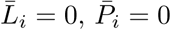 for all phage variants *i*. This equilibrium is unstable as long as the bacterial logistic growth rate *r >* 0.

Secondly, there is an equilibrium in which the susceptible population is at carrying capacity and the infection is absent: 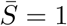, and 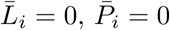 for all phage variants *i*. This equilibrium is stable if no phage-lysogen pairs *P*_*i*_-*L*_*i*_ can invade on the susceptible population. To derive stability conditions, we approximate the dynamics of *P*_*i*_ and *L*_*i*_ in the vicinity of the equilibrium by the linearised equations

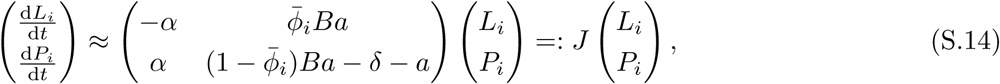

where 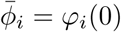 is the lysogeny propensity of phage type *i* at the equilibrium. No phage-lysogen pair can invade (*i*.*e*., the equilibrium is stable) precisely if the real parts of both eigenvalues of the Jacobian matrix *J* are negative for all *i*. The eigenvalues of the Jacobian matrix *J* are given by

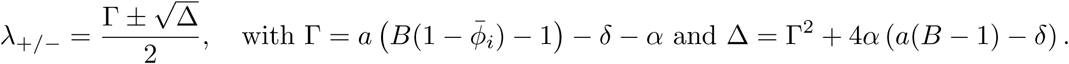

The real parts of *λ*_+*/−*_ are both negative precisely if Δ *<* Γ^2^ **and** Γ *<* 0. From Γ *<* 0, we find

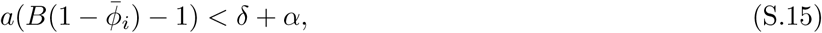

while Δ *<* Γ^2^ yields

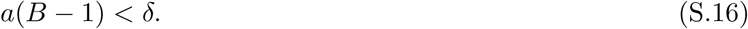

Since all parameters are non-negative and 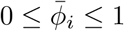, condition S.16 is more stringent than condition S.15. Note also that condition S.16 does not depend on the lysogeny propensity of the invading phages, 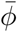. Hence, the equilibrium 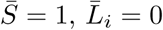, and 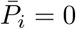 for all phage variants *i* is stable exactly if condition S.16 is satisfied. This condition makes sense: phages cannot spread in a susceptible cell population at carrying capacity if their infection rate 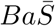 is smaller than the decay rate of phage particles 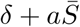.

Thirdly, there is a class of equilibria in which 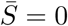, and 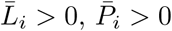 for some *i*. In these equilibria, the total densities of lysogens 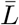 and of free phages 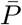 are given by

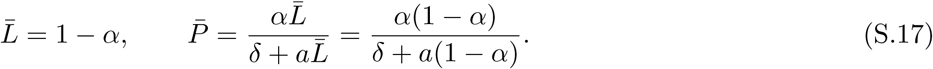

In the absence of susceptible cells at equilibrium, no infections can take place and hence all phage variants behave identically (since phages vary only in *φ*_*i*_(*A*), which occurs exclusively in the infection terms). This is reflected in the equations for the different lysogen variants, which for 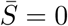 and 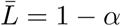 (Eq S.17) reduce to

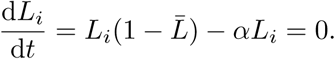

Hence, any combination of 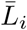 values with 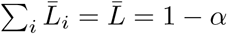 and corresponding 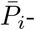values,

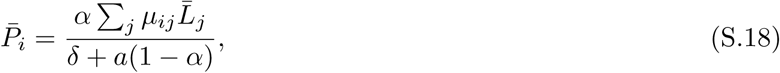

is an equilibrium. Analogous to the reasoning above, such an equilibrium is stable if the susceptible cells cannot invade the phage-lysogen population at equilibrium. The linearised equation for the dynamics of *S* near the equilibrium of Eq S.17 reads

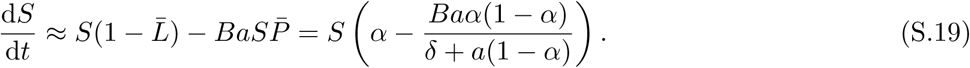

The right-hand side of this equation is negative (*i*.*e*., susceptible cells cannot invade) precisely if *δ* + *a*(1 − *α*) *< Ba*(1 − *α*), or summarised

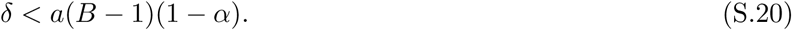

(Note that to arrive at condition (S.20) we assume *δ* + *a*(1 − *α*) *>* 0. This assumption is justified because the spontaneous induction rate of lysogens *α* is small, and hence 1 − *α >* 0 (see Supplementary Table 1).)

Lastly, there can be an equilibrium in which the susceptible cells, lysogens and phages all coexist. Expressions for 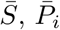 and 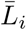 at this equilibrium are bulky and not directly insightful. This type of equilibrium was however extensively analysed in recent work by Wahl *et al*. for phages with a constant lysogeny propensity (*i*.*e*., the restricted model where arbitrium communication is excluded) [16]. Remember that in an equilibrium state, phage variants are characterised by their lysogeny propensity *φ*_*i*_(*Ā*) only (*i*.*e*., differences in response threshold *θ* are relevant only if they are reflected in differences in *φ*_*i*_(*Ā*); phage variants *i* and *j* with *θ*_*i*_ ≠ *θ*_*j*_ but *φ*_*i*_(*Ā*) = *φ*_*j*_(*Ā*) can for all practical purposes be considered the same), and hence the results found by Wahl *et al*. can be extended to the model analysed here. Phage variants with different lysogeny propensity *φ*_*i*_(*Ā*) can be seen as consumers that compete for a single resource, namely susceptible cells to infect. As long as 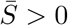, we hence expect competitive exclusion, and the phage population at equilibrium will be dominated by phages with a single lysogeny propensity value 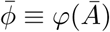 (when mutations are ignored, these phages will be the only ones present; otherwise we find a quasispecies). Furthermore, Wahl *et al*. show that if susceptible host cells coexist with a resident lysogen-phage population with some lysogeny propensity 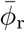, a phage-lysogen pair of a variant with higher lysogeny propensity 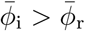 can always invade on this equilibrium. Hence, in this equilibrium, the dominant phage will be the one with the highest equilibrium lysogeny propensity, 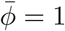.

As has been demonstrated previously [11, 16], the “coexistence equilibrium” is stable only if phages and lysogens can invade on a susceptible population at carrying capacity (*i*.*e*., condition S.16 is violated), and susceptible cells can invade on the phage-lysogen population in equilibrium (*i*.*e*., condition S.20 is violated). Hence, susceptible cells, lysogens and phages all coexist precisely if

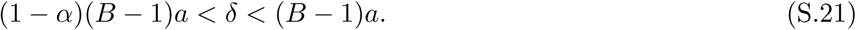

If the effective burst size *B >* 1 (a necessary condition for the phage to be viable), this can be rewritten as

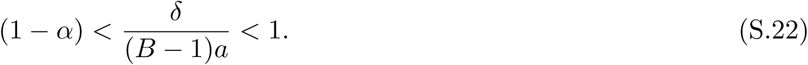

Because the spontaneous induction rate of lysogens, *α*, is small (*α <* 0.01, see Supplementary Table 1), condition S.22 is very specific. Susceptible cells, lysogens and phages coexist only if the exponential growth rate of a lytic phage spreading in a susceptible population at carrying capacity, (*B* − 1)*a −δ*, is positive but very small, *i*.*e*., if the epidemic is viable but only barely so. In reality, however, most phage epidemics are characterised by a high infectivity, mainly because of a large burst size [31]. Therefore, condition S.22 is rarely satisfied, and for most phages we should instead expect to converge to equilibria of the third type 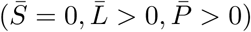.

This observation has consequences for the selection pressures on phage variants over the course of a typical epidemic. As soon as the pool of susceptible host cells is depleted, competition between the different phage variants vanishes and the relative frequency of the variants freezes (see Supplementary Fig S1 for an illustration). Under these conditions, no infections take place and hence there is no selection on the lysis-lysogeny decision. We conclude that the evolution of the lysis-lysogeny decision of typical phages requires regular perturbations away from equilibrium conditions.

### S3 Derivation of the Evolutionarily Stable Strategy (ESS) under the serial-passaging regime

In the main text, we present simulation results of the evolution of the lysis-lysogeny decision of phages exposed to a serial-passaging regime, both when arbitrium communication is excluded from the model (Fig 2), and when it is included (Fig 3, 4 and Supplementary Fig S2). These simulations show that, after many passaging episodes, a single variant dominates the phage population (accompanied by its quasispecies, see Fig 2C, Fig 3B, and Supplementary Fig S2). We therefore assume that, within the parameter range of interest, a pure Evolutionary Stable Strategy (ESS) exists. In this section, we derive analytical expressions for the ESS. We first provide a definition of the ESS under a serial-passaging regime, and give a general description of how this ESS can be found (section S3.1). Then, we apply this general approach to derive the ESS of phages that differ in a constant lysogeny propensity, *ϕ* (absence of arbitrium communication, section S3.2), and the ESS of communicating phages that differ in their response threshold *θ* (section S3.3).

#### S3.1 General approach

Consider a population of phages under a serial-passaging regime with long time *T* between passages. At the start of each passaging episode, a fraction *D* of the phages is taken from the end of the previous episode, and is added to a “fresh” population of susceptible bacteria at carrying capacity. This procedure is repeated over many episodes. Within each episode, the dynamics of the susceptible bacteria, lysogens, phages, and arbitrium are described by Eq S.7–S.12.

An **evolutionarily stable strategy (ESS)** is defined as a strategy that cannot be invaded by any other strategy that is initially rare. To find the ESS, we therefore consider a scenario where an invader phage variant attempts to invade a resident phage variant. Below, we specify what it means for a phage variant to be able to invade in a resident phage population under the imposed serial-passaging regime.

Envision a resident population consisting of an isogenic phage population that has gone through many passaging episodes. Over time, the dynamics within these episodes have converged to a repeatable trajectory characterised by *P*_r_(*t*), the resident phage density over time, *S*(*t*), the density of susceptible bacteria over time, and *L*_r_(*t*), the density of lysogens over time. At the start of one episode, now suddenly introduce a second phage with its own (possibly different) strategy. Crucially, the initial density of this invading phage *P*_i_(0) is infinitesimally small. Consequently, during the first passaging episode the dynamics of the resident phage and the bacteria (*P*_r_(*t*), *S*(*t*), and *L*_r_(*t*)) are not affected by the invader. The invader is able to invade precisely if at the end of the first episode its frequency has increased relative to that of the resident, *i*.*e*., if *P*_i_(*T*)*/P*_r_(*T*) *> P*_i_(0)*/P*_r_(0).

Note that the relative frequency of an invader with exactly the same strategy as the resident does not change during an episode. Suppose that such an invader is the best-performing invader under the environment set up by the resident; then this implies that no invader can increase in frequency over a passaging episode, and therefore the resident strategy must be an ESS. In other words, in a resident phage population consisting of the ESS only, the ESS itself is the optimal strategy for an invading phage variant, *i*.*e*., *the ESS is the optimal response to itself*.

What does it take to be the best-performing invader? To answer this question, we consider the dynamics within a single passaging episode in more detail.

If the time between passages *T* is long, and the parameter conditions are such that the system converges to an equilibrium with *S* = 0, *P >* 0, *L >* 0 (typical parameter values, see sections S1.4 and S2), the dynamics within an episode can be divided in three distinct phases (see Supplementary Fig S1):

1. **Epidemic phase:** A substantial population of susceptible bacteria (*S >* 0) supports an ongoing epidemic. Free phages and lysogens are formed through infection of susceptible bacteria.
2. **Transition phase:** The population of susceptible cells has collapsed (*S ≈* 0). The lysogen population expands to fill up the space left behind by lysed cells. Free phages particles can no longer infect susceptible cells, and disappear through decay and adsorption to lysogens.
3. **Equilibrium phase:**The composition of the population is well-characterised by an equilibrium of “type 3” (see Eq S.17). There is a small but consistent population of free phages that originates from lysogens through spontaneous induction. The distribution of phage variants in the free phage population is a direct representation of the relative frequency of the variants in the lysogen population (Eq S.18).

Let *T*_E_ be the time at which the susceptible population collapses, *i*.*e*., the end of the epidemic phase. If the time between passages *T* is sufficiently larger than *T*_E_, the passage takes place during the equilibrium phase. The relative frequency of phage variants in the transferred sample then directly reflects the relative frequency of the corresponding lysogens. Since lysogens are only differentially formed through infection dynamics (and not through lysogen replication, which happens at the same rate for all lysogen variants), the relative frequency of the different lysogens is established during the epidemic phase and does not change afterwards. The performance of an invading phage can hence be measured by its lysogen production between *t* = 0 and *t* = *T*_E_.

The dynamics of *S*(*t*), and consequently *T*_E_, are determined by the resident phage: highly virulent resident phages (that cause many lytic infections, for instance because of a low *ϕ*-value) deplete the susceptible cell population faster than less virulent residents. The optimal invader under a certain resident is the phage variant that produces the most lysogens during the limited window of opportunity that it is offered by the environment set up by the resident. Since the ESS is the optimal response to itself, it is the strategy that, as an invader, produces the most lysogens during an epidemic phase set up by itself. We will use this reasoning to identify the ESS.

#### S3.2 Evolutionarily stable lysogeny propensity *ϕ*^*∗*^ of non-communicating phages

Consider the restricted model in which arbitrium communication is excluded and phages are characterised by a fixed lysogeny propensity *ϕ*. To find the ESS under this scenario, we take the following steps:

1. Derive how the duration of the epidemic, *T*_E_, depends on the *ϕ*-value of a resident phage population.
2. Find the optimal *ϕ* given a fixed value of *T*_E_, *i*.*e*., the *ϕ*-value that yields a maximal lysogen density at time *T*_E_.
3. Combine 1. and 2. to find the ESS: the *ϕ*-value *ϕ*^*∗*^ that maximises its lysogen density at time *T*_E_(*ϕ*^*∗*^), the duration of the epidemic as dictated by its own *ϕ*-value, *ϕ*^*∗*^.

##### S3.2.1 Simplifying assumptions

To make the model analytically tractable, we make the following simplifying assumptions (based on the typical infection dynamics, see Fig 2A and Supplementary Figure S1):

1. Bacterial growth, decay of free phages and induction of lysogens are considered to be much slower than the phage infection dynamics. We hence ignore these processes when describing the epidemic phase.
2. The epidemic ends when all susceptible cells have been infected. In other words, we solve *T*_E_ from 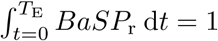 (where this 1 represents the carrying capacity) in the non-dimensionalised units).
3. The density of lysogens during the epidemic is small, hence *N ≈ S*.
4. The susceptible population remains relatively constant for some time, after which it rapidly collapses. We approximate these dynamics with a block function, setting *S* = 1 for *t ≤ T*_E_, and *S* = 0 for *t > T*_E_.

Under these assumptions the dynamics of the resident phage population for the period 0 *≤ t ≤ T*_E_ are described by:

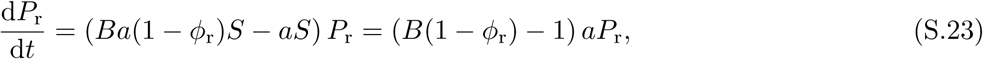

with solution

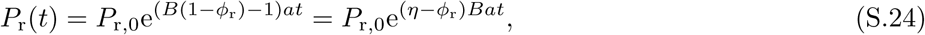

where *P*_r,0_ *≡ P*_r_(0) is the initial density of resident phages and we have introduced *η* := 1 − *B*^−1^. Note that the description of the infection dynamics in Eq S.23 is meaningful only if early in the epidemic the phage population indeed grows exponentially, *i.e.*, if *ϕ*_r_ *< η*. For default parameter settings, this upper bound on *ϕ*_r_ is well above the *ϕ*-values that are typically selected (*ϕ ≈* 0.04 (Fig 2C), while *η* = 0.5 (Supplementary Table 1)), indicating that this assumption is reasonable.

##### S3.2.2 Duration of the epidemic *T*_E_ given a resident phage

First, we derive how the duration of the epidemic, *T*_E_, depends on the lysogeny propensity of a resident phage variant *ϕ*_r_. To find *T*_E_(*ϕ*_r_), we substitute Eq S.24 into assumption 2:

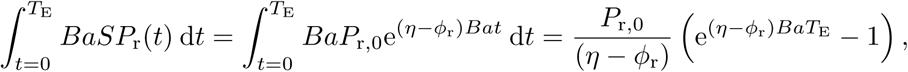

and then equate this integral to 1 to find

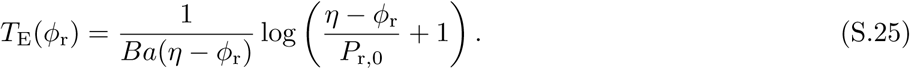

Note that the density of the resident phage at time *T*_E_ is now given by

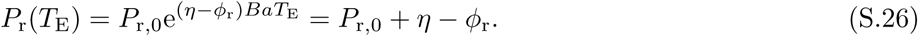

Therefore, the expression for *T*_E_ (Eq S.25) can also be read as:

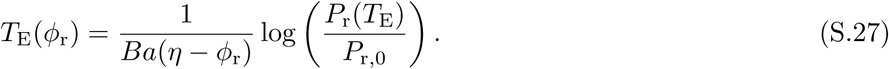

Eq S.27 will prove useful later for the derivation of the ESS.

##### S3.2.3 Optimal invader strategy *ϕ*_i_ given a fixed duration of the epidemic *T*_E_

Next, we ask what invader lysogen propensity *ϕ*_i,opt_ maximises the invader’s lysogen production, *L*_i_(*T*_E_), if the duration of the epidemic *T*_E_ is fixed. For 0 *≤ t ≤ T*_E_, the dynamics of *L*_*i*_(*t*) are described by

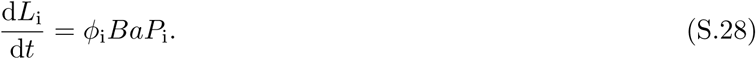

Since 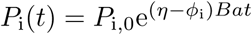 (see Eq S.24) and *L*_i_(0) = 0, we can now solve

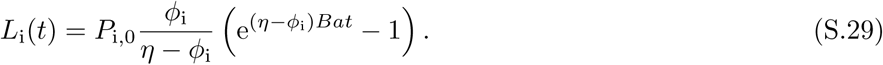

To find the *ϕ*_i_-value that maximises *L*_i_(*T*_E_), we take the derivative of Eq S.29 with respect to *ϕ*_i_:

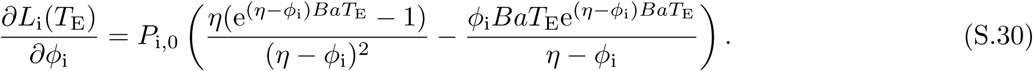

To find *ϕ*_i,opt_, we should hence solve

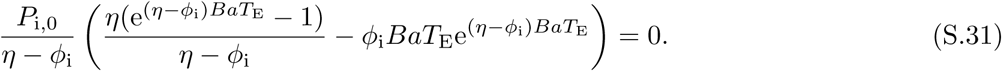

Unfortunately, Eq S.31 cannot be solved analytically. We can however simplify Eq S.31 by noting that for sufficiently small *ϕ*_i_, (*η* − *ϕ*_i_) is of order 0.1 − 1, while *T*_E_ is typically of order 1 (see Supplementary Figure 1), and *Ba* is of order 10 − 1000 (Supplementary Table 1). Hence, 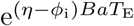 is typically ≫ 1, and Eq S.31 can be approximated by

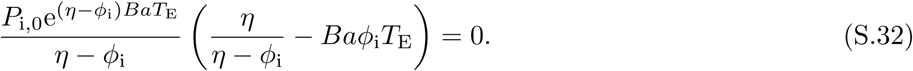

From Eq S.32 we find

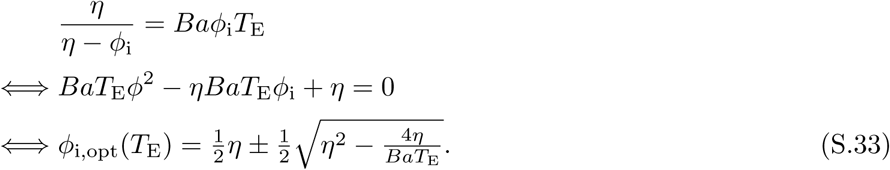

##### S3.2.4 The ESS

Eq S.25 and its alternative formulation Eq S.27 describe how the duration of the epidemic *T*_E_ depends on the lysogeny propensity *ϕ*_r_ of the current resident, while Eq S.33 gives the value *ϕ*_i,opt_ that maximises the lysogen production of an invader during an epidemic of a fixed duration *T*_E_. The ESS is now given by the value *ϕ*^*∗*^ that is “optimal” as defined by Eq S.33, when it itself is the resident and hence dictates *T*_E_(*ϕ*^*∗*^). Combining Eq S.33 and Eq S.27 we find

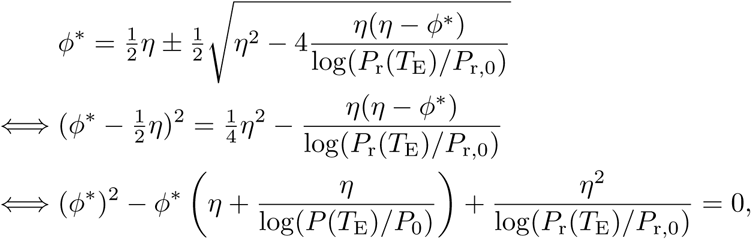

from which we can solve:

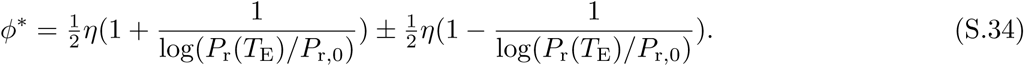

Of these two solutions, *ϕ*_+_ = *η* is an asymptote at which our approximation no longer holds (remember that we previously demanded that *ϕ*_r_ *< η* to ensure initial spread of the infection). Hence, *ϕ*^*∗*^ should be given by *ϕ*_−_:

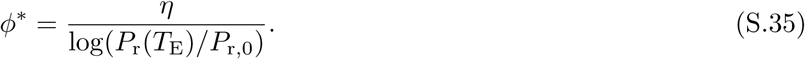

Although Eq S.35 seems to provide an elegant equation for the ESS, it still depends on *P*_r_(*T*_E_) and *P*_r,0_. If the interval between passages is sufficiently long, the phage density at the end of a passaging cycle will be given by Eq S.17 and hence

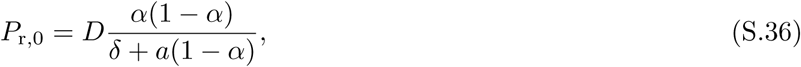

where *D* is the dilution factor of phages upon passaging. While the value of *P*_r,0_ does not depend on the lysogeny propensity *ϕ*_r_, *P*_r_(*T*_E_) does (see Eq S.26). Substituting *P*_r_(*T*_E_) = *P*_r,0_ + *η* − *ϕ*^*∗*^ yields

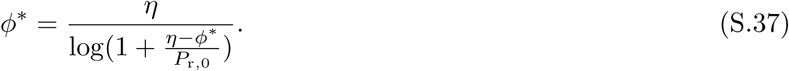

This equation cannot be solved analytically. However, we can make a reasonable approximation of Eq S.37 by considering the differences in orders of magnitude of the terms within the logarithm. As argued above, (*η* − *ϕ*^*∗*^) is generally of order 0.1 − 1, while typical values of *P*_r,0_ are several orders of magnitude smaller (*P*_r,0_ *≈* 10^−5^). Therefore, we can approximate the logarithm in Eq S.37 by

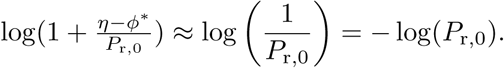

Using this approximation, we find

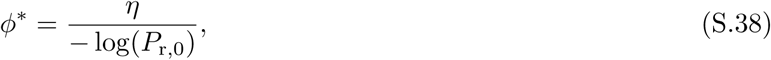

which is also presented in the main text (Eq 5). Eq S.38 and S.36 were used to find the analytical predictions shown in Fig 2D.

#### S3.3 Evolutionarily stable response threshold *θ*^*∗*^ of communicating phages

Next, consider a population of phages that do engage in arbitrium communication, again under a serial-passaging regime with sufficiently long time between the passages. Below, we use an approach similar to section S3.2, but more general, to derive the evolutionarily stable arbitrium response threshold, *θ*^*∗*^. We take the following steps:

1. Describe the dynamics of an invading phage and its corresponding lysogens in an environment dictated by a resident phage.
2. Find the optimal invader response threshold under a fixed resident response threshold, *i*.*e*., find the *θ*-value that maximises the invader’s lysogen production at time *T*_E_ when the dynamics of susceptible cells (and hence *T*_E_) are determined by a fixed resident phage.
3. Determine the ESS, *θ*^*∗*^, as the optimal response to itself: the optimal invader response threshold (as found in step 2) if that same response threshold is the resident strategy.

We found that the results below are best understood in terms of the non-scaled model; in particular the (non-scaled) burst size of the phages turns out to be an important parameter. Therefore, the derivations below are presented for the dimensionalised equations Eq S.1–S.6.

##### S3.3.1 Simplifying assumptions

To make the model tractable, we again make a few simplifying assumptions:

1. As in section S3.2, we assume that there is a separation of time scales between the infection dynamics of the phages and the reproduction of the bacteria, spontaneous phage decay and lysogen induction. Hence, when describing the epidemic phase we ignore these other processes.
2. Additionally, we assume that there also is a separation of time scales between the production of arbitrium through infections (first term in Eq S.4) and its uptake and degradation by cells (second term in Eq S.4). We ignore the uptake and degradation of arbitrium during the early epidemic, such that the increasing arbitrium concentration reflects the decrease of the susceptible cell density because of infections.
3. We assume that communicating phages switch from a completely lytic strategy (*φ*(*A*) = 0) to a completely lysogenic strategy (*φ*(*A*) = 1) once the arbitrium concentration exceeds the phages’ response threshold. This is in line with observations from simulations, where we find that phage variants with *ϕ*_max_ = 1 dominate the population for a wide range of parameter values (Fig 4A).

The assumptions above are less strict then the assumptions made in section S3.2. In particular, we no longer assume that the density of susceptible cells, *S*(*t*), remains constant for the duration of the epidemic 0 *≤ t ≤ T*_E_. Rather, for the derivations below it suffices to assume that *S*(*t*) is a declining function which is completely determined by the resident phage, and that *S*(*t*) is sufficiently close to 0 after the epidemic, *i*.*e*., for times *t > T*_E_ (where *T*_E_ still depends on the characteristics of the resident phage).

It will be useful to refer to the *time* when the resident and invader switch to lysogeny as *τ*_r_ and *τ*_i_, respectively. Given particular dynamics of the susceptible cell density and the arbitrium concentration dictated by a resident phage population, there is a direct relation between *τ*_i_ and *θ*_i_ (the arbitrium response threshold of the invader). Keep in mind, however, that this relation changes if the resident phage is changed.

##### S3.3.2 Dynamics of the invading phage and its corresponding lysogens

First, we describe how the dynamics of an invading phage variant depend on the resident phage population. Remember that we consider an invader phage variant that starts off at infinitesimally small density, and attempts to invade an isogenic resident phage population that has already converged to a repeatable trajectory of *P*_r_(*t*), *L*_r_(*t*), *S*(*t*), and *A*(*t*) per passaging episode. Under these conditions, the dynamics of *S*(*t*), *N* (*t*) = *S*(*t*) + *L*_r_(*t*) and *A*(*t*) over the first passaging episode do not depend on the switch time *τ*_i_ of the invader, but only on the switch time of the resident phage, *τ*_r_. Based on the assumptions formulated above, the ODEs for the density of the invading phage and its corresponding lysogens can be written as

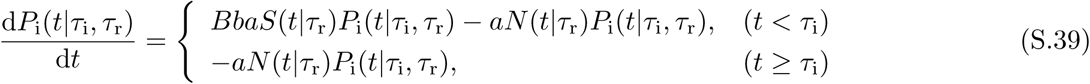

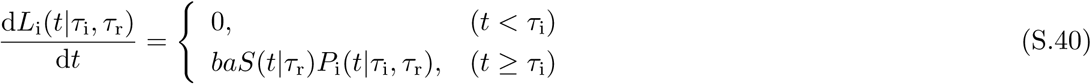

where vertical lines are used to indicate which phage charactistics the trajectories of variables depend upon. Eq S.39–S.40 capture the switch from a completely lytic infection strategy (for *t ≤ τ*_i_), in which new phage particles are produced through infection but no lysogens are formed, to a completely lysogenic strategy (for *t > τ*_i_), in which no new phage particles are produced but all infections result in the production of lysogens (see assumption 3). Remember that we here use the dimensionalised equations, so *B* is the burst size, *a* the rate of adsorption of phages to bacterial cells (irrespective of whether they are susceptible or lysogen), and *b* the probability that adsorption to a susceptible cell leads to an infection.

The solution to Eq S.39 can be written as

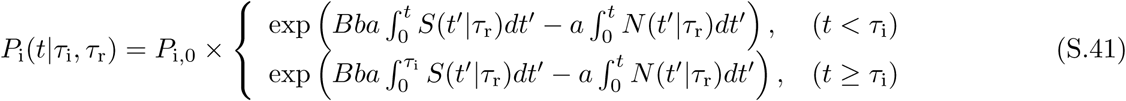

as is easily verified by differentiating this solution with respect to *t*.

As before, the performance of the invading phage variant is determined by its lysogen production during the epidemic phase, *i*.*e*., between *t* = 0 and *t* = *T*_E_. At any time *t*, the density of invader lysogens is

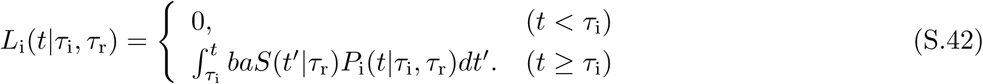

Invading phage variants are selected on their lysogen density at the end of the epidemic, *L*_i_(*T*_E_|*τ*_i_, *τ*_r_). Once the epidemic phase has ended (*t ≥ T*_E_), no new phage particles or lysogens are formed through infection. Hence, any reasonable switching time must obey *τ*_i_ *< T*_E_. Furthermore, since *S*(*t*) *≈* 0 for any time *t ≥ T*_E_,

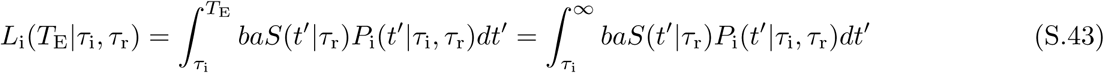

which we will denote Λ_i_(*τ*_i_, *τ*_r_).

##### S3.3.3 Optimal invader strategy *τ*_i_ given some resident phage

The optimal invader strategy *τ*_i_ given *S*(*t*|*τ*_r_) and *N* (*t*|*τ*_r_) is the one that maximises Λ_i_(*τ*_i_, *τ*_r_). To find this optimal strategy, we differentiate Eq S.43 with respect to *τ*_i_:

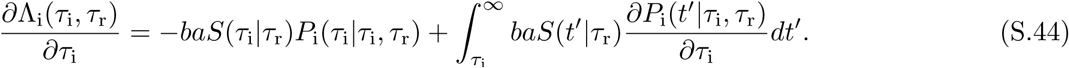

The derivative in the integrand can be calculated from Eq S.41 (noting that, inside the integral, *t′ ≥ τ*_i_):

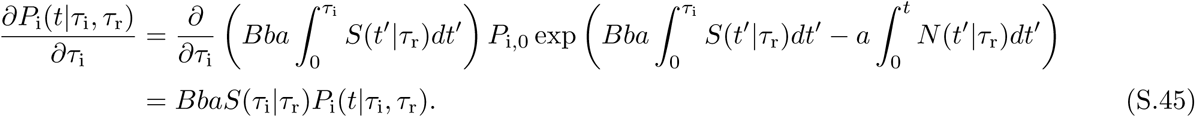

Inserting the last expression into Eq S.44 yields

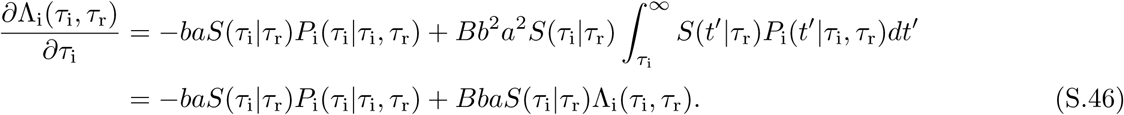

The terms in Eq S.46 have a clear interpretation. By taking the derivative of Λ_i_(*τ*_i_, *τ*_r_) to *τ*_i_, we are implicitly comparing one possible invading phage variant (phage 1) that switches at time *t* = *τ*_i_ to a second invading phage variant (phage 2) that switches ever so slightly later, at *t* = *τ*_i_ + d*τ*. Eq S.46 says that the lysogen density of these two variants at the end of the epidemic will differ because of two effects: On the one hand (first term) phage 2 will have a *lower* lysogen density than phage 1 because it does not produce lysogens in the time interval from *τ*_i_ to *τ*_i_ + d*τ*. The damage is *−baS*(*τ*_i_|*τ*_r_)*P*_i_(*τ*_i_|*τ*_i_, *τ*_r_)d*τ* lysogens per volume. On the other hand, phage 2 will have a higher *higher* lysogen density because it produces additional free phages in the time interval from *τ* to *τ* + d*τ*, which results in additional lysogens in the rest of the epidemic. As a result, throughout the rest of the epidemic the second phage has (1 + *BbaS*(*τ*_i_|*τ*_r_)d*τ*) times as many phages as the first phage variant, and therefore produces an additional number of *BbaS*(*τ*_i_|*τ*_r_)*L*_i_(*τ*_i_, *τ*_r_)d*τ* lysogens per volume.

The optimal invading phage variant given a resident phage is the variant with the value of *τ*_i,opt_(*τ*_r_) for which the two terms in Eq S.46 cancel precisely:

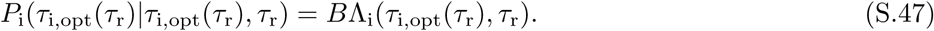

That is, the optimal invader switches precisely when its phage density is equal to its total eventual lysogen production multiplied by the burst size *B*.

We may rewrite Eq S.47 as

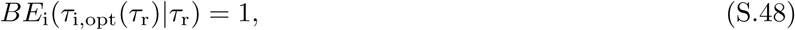

where *E*_i_(*τ*_i_|*τ*_r_) *≡* Λ_i_(*τ*_i_, *τ*_r_)*/P*_i_(*τ*_i_|*τ*_i_, *τ*_r_) is the number of lysogens eventually produced per phage of the invader phage variant present at time *τ*_i_. *E*_i_(*τ*_i_|*τ*_r_) can be interpreted as a kind of “exchange rate”, expressing the value of a single phage at time *τ*_i_ in the currency of lysogens. This suggests another way of phrasing the results above, where we compared two phage variants of which phage 2 switched slightly later than phage 1: During the time interval from *τ*_i,1_ to *τ*_i,2_ = *τ*_i,1_ + d*τ*, both competing invading phage variants infect *baS*(*τ*_i,1_)*P*_i_(*τ*_i,1_|*τ*_i,1_, *τ*_r_)d*τ* susceptible bacteria per volume. Phage 1 directly converts these infected bacteria into lysogens. Phage 2 instead converts each of them into *B* additional phages. Whether this is a good idea depends precisely on whether increasing the phage density by *B* phages per volume will, during the rest of the epidemic, result in an increased lysogen density of more than 1 lysogen per volume. That is, phage 2 is the better invader precisely if *BE*_i_(*τ*_i_|*τ*_r_) *>* 1, while phage 1 is the better invader if *BE*_i_(*τ*_i_|*τ*_r_) *<* 1. Again we see that the optimal invader must obey Eq S.47 and S.48.

##### S3.3.4 The ESS

To find the ESS, we ask what phage variant is the optimal response to itself, *i*.*e*., what phage variant satisfies

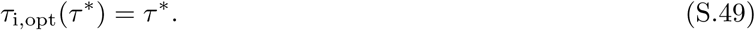

In other words, the ESS must obey a special case of Eq S.47 and S.48:

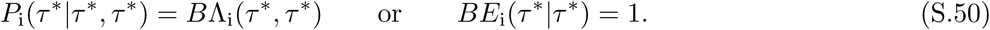

Importantly, in this case the resident and invader behave identically, so that the exchange rate of the resident must be the same as that of the invader. That is,

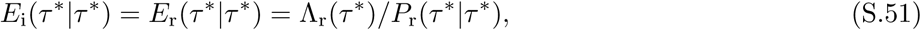

where Λ_r_(*τ*_r_) is the total density of lysogens eventually produced by a resident with switching time *τ*_r_. Combined with Eq S.50 this results in:

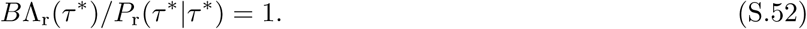

Hence, the ESS is the strategy that, when it is the only phage variant present, switches from the lytic to the lysogenic cycle precisely when the density of free phage particles it has is equal to the burst size times the density of lysogens it will still produce in the remainder of the active epidemic (see explanation in the previous section).

So far, we have expressed the results for the ESS as a switching time *τ*^*∗*^. In reality, however, the communicating phages switch when a certain threshold arbitrium concentration *θ*^*∗*^ is reached. As a last step, we therefore have to relate the terms in Eq S.52 to the arbitrium concentration. Under our simplifying assumptions, the arbitrium dynamics between time *t* = 0 and *t* = *τ*_r_ are described by

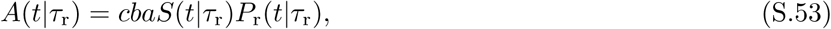

where *c* is increase in arbitrium concentration per infection. The total arbitrium concentration at time *t* is hence given by

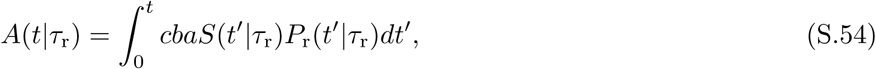

which can be written as *A*(*t*|*τ*_r_) = *cI*_r_(*t*|*τ*_r_), where

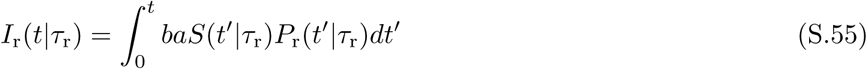

is the infection density: the number of infections that has occurred per volume at time *t*.

To express the ESS in terms of the arbitrium concentration, we first show that *P*_r_(*τ*^*∗*^|*τ*^*∗*^) is approximately proportional to *I*_r_(*τ*^*∗*^|*τ*^*∗*^). In general, the resident phage density obeys an equation equivalent to Eq S.39 (even though this equation was originally written down for the invading phage). For the time period *t < τ*_r_, the solution of this equation can be expressed as

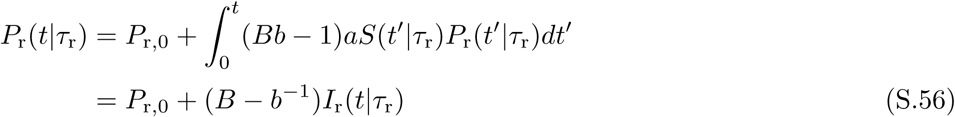

Provided that the initial phage density *P*_r,0_ is negligible compared to the phage density at time *τ*^*∗*^, we find that

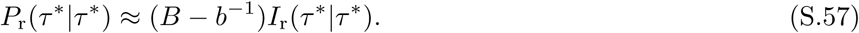

Next, we use that the epidemic will eventually consume (almost) all susceptible bacteria. (Note that this is equivalent with our earlier assumption that *S*(*t*) *≈* 0 for *t > T*_E_.) Hence, we must have that

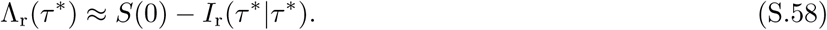

If we insert Eq S.57 and S.58 into Eq S.47 and solve for *I*_r_(*τ*^*∗*^|*τ*^*∗*^), we arrive at

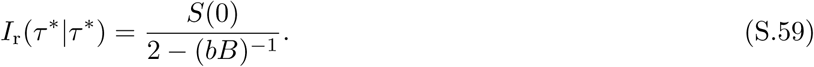

That is, the ESS switches when the infection density obeys Eq S.59. This implies that the ESS should have the threshold

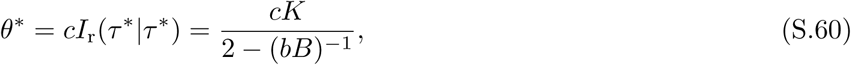

where we have substituted *S*(0) = *K*, the carrying capacity of the bacteria. Eq S.60 is also presented in the main text (Eq 6). This equation was used to provide the analytical estimates shown in Fig 4B.

### S4 Supplementary Figures

**Supplementary Figure S1.**
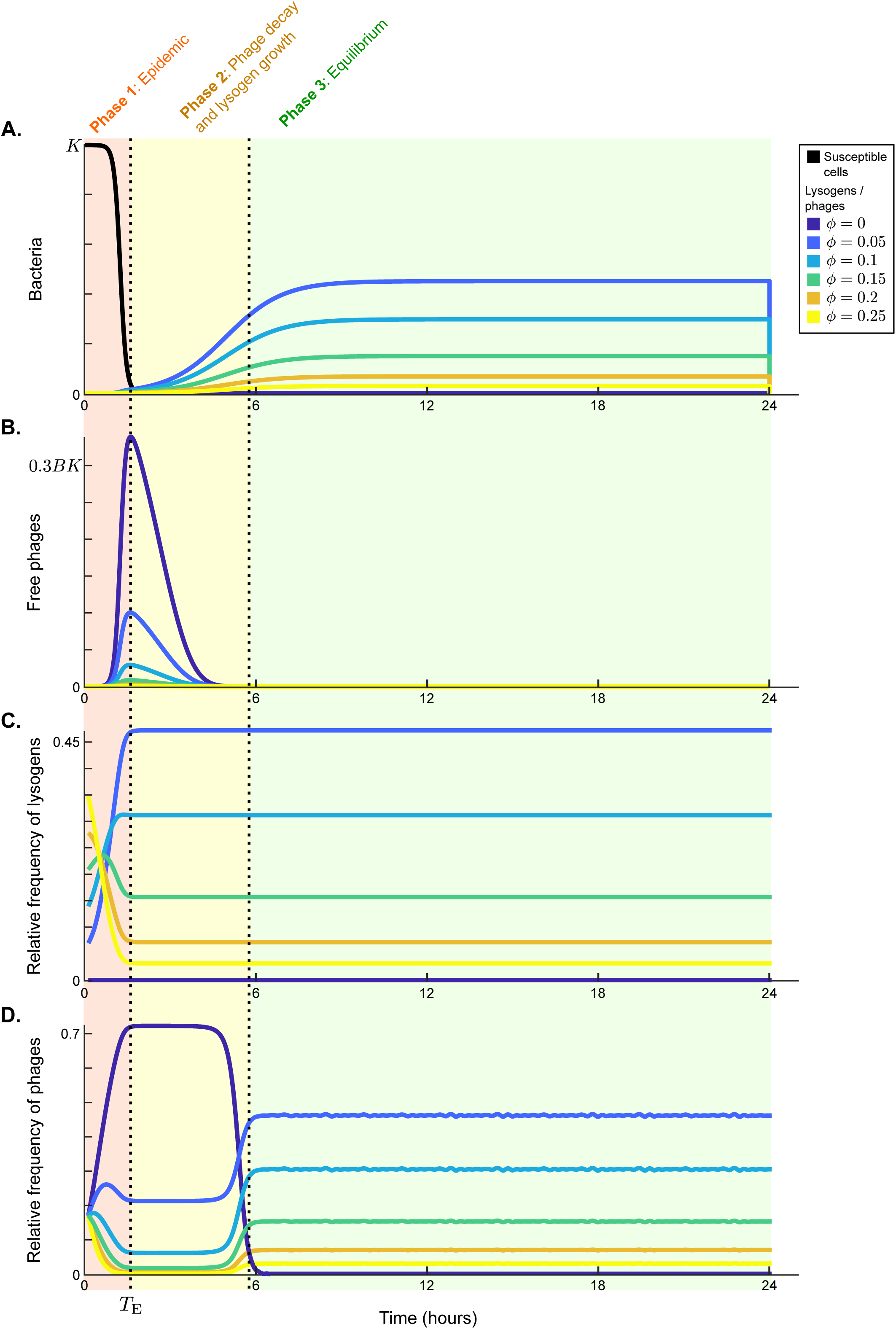
Model dynamics during a single passaging episode for default parameter values and constant lysogeny propensities (no communication). Absolute frequencies of susceptible cells and lysogens (panel A) and phages (panel B) are plotted as a function of time, as well as the relative frequency of the different phage variants in the lysogen population (panel C) and the free phage population (panel D). The dynamics can be divided into three distinct phases: (1) Epidemic: Susceptible cells are still present (*S >* 0) but are being depleted by infection. Phage and lysogen densities increase depending on their lysogeny propensity *ϕ*_*i*_. The relative frequencies in the lysogen population are established. (2) Transition phase: Infections no longer take place because the susceptible cell population has been depleted (*S ≈* 0). The lysogen population grows until it reaches its carrying capacity. Free phage densities decline because of natural decay and adsorption to lysogens. The influx of free phages by reactivation of lysogens differs per phage variant because the density of their corresponding lysogens differs, and the relative frequencies in the free phage pool change to reflect this. (3) Equilibrium phase: The total lysogen population is at carrying capacity, and the relative frequencies of the phage variants in the (eventually passaged) free phage pool reflect the relative frequencies of phage variants in the lysogen population.

**Supplementary Figure S2.**
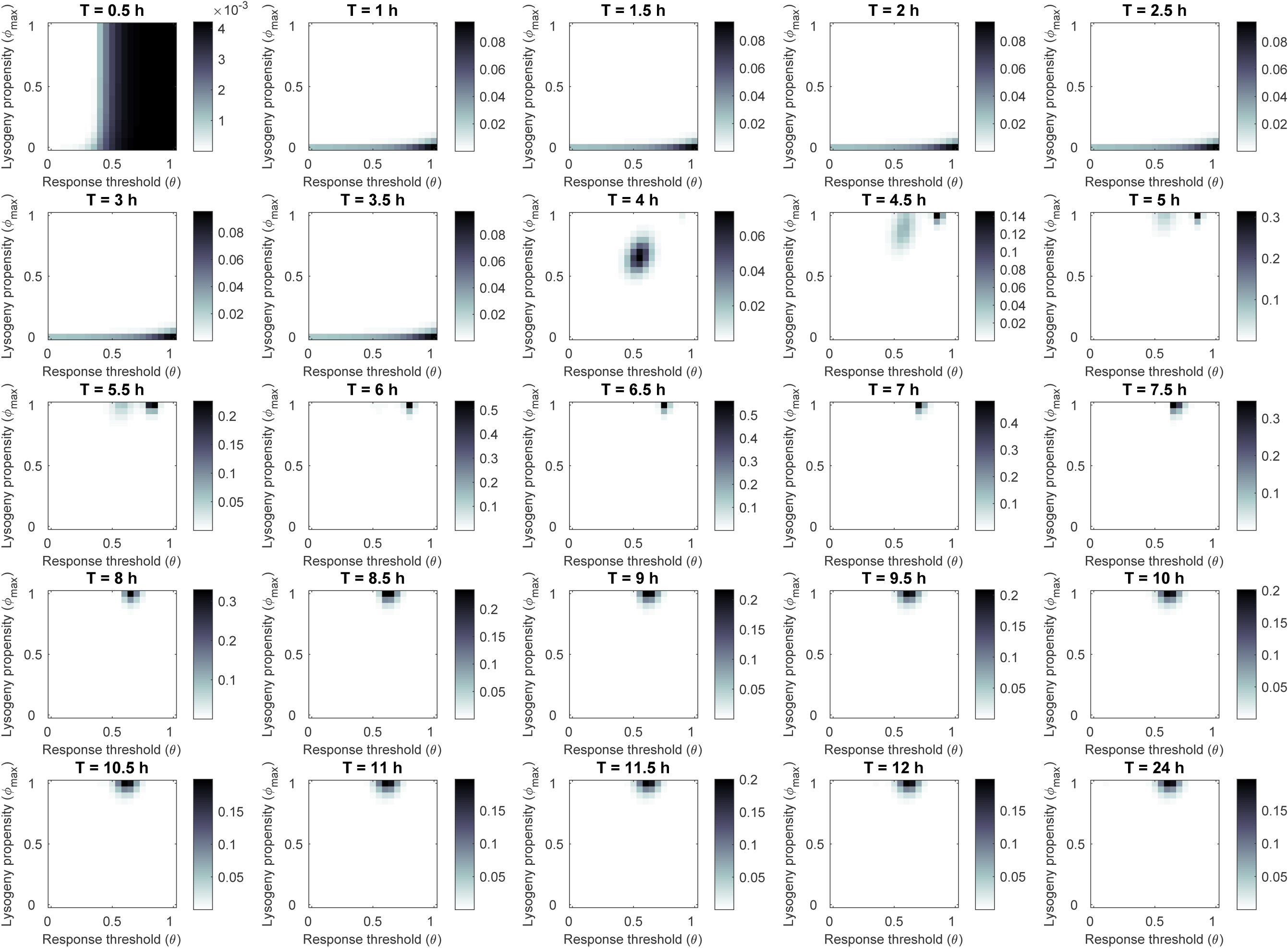
Most abundant phage variant at evolutionary steady state for different lengths of the time interval between passages *T*. Default parameter settings (Supplementary Table S1). In each run, 441 variants were included: all combinations of *ϕ*_max_ = 0, 0.05, *…*, 1 and *θ* = 0, 0.05*cK, …, cK*. Simulations were run for 1000 passaging episodes to reach evolutionary steady state, after which the phage variant distribution was plotted. For *T <* 4 h phages with a fully lytic strategy (*ϕ*_max_ = 0) are selected, while for *T ≥* 5 h phages are selected that switch from a fully lytic strategy at low arbitrium concentration to a fully lysogenic strategy (*ϕ*_max_ = 1) at high arbitrium concentration.

## References

1. Nealson KH, Platt T, Hastings JW. Cellular Control of the Synthesis and Activity of the Bacterial Luminescent System1. Journal of Bacteriology. 1970;104(1):313–322.

2. Miller MB, Bassler BL. Quorum Sensing in Bacteria. Annual Review of Microbiology. 2001;55(1):165–199. doi: 10.1146/annurev.micro.55.1.165.

3. Hense BA, Schuster M. Core Principles of Bacterial Autoinducer Systems. Microbiol Mol Biol Rev. 2015;79(1):153–169. doi: 10.1128/MMBR.00024-14.

4. Antunes LCM, Ferreira RBR, Buckner MMC, Finlay BB. Quorum sensing in bacterial virulence. Microbiology (Reading, England). 2010;156(Pt 8):2271–2282. doi: 10.1099/mic.0.038794-0.

5. Diggle SP, Griffin AS, Campbell GS, West SA. Cooperation and conflict in quorum-sensing bacterial populations. Nature. 2007;450(7168):411–414. doi: 10.1038/nature06279.

6. Darch SE, West SA, Winzer K, Diggle SP. Density-dependent fitness benefits in quorum-sensing bacterial populations. Proceedings of the National Academy of Sciences. 2012;109(21):8259–8263. doi: 10.1073/pnas.1118131109.

7. Cornforth DM, Foster KR. Competition sensing: the social side of bacterial stress responses. Nature Reviews Microbiology. 2013;11(4):285–293. doi: 10.1038/nrmicro2977.

8. Kleerebezem M, Quadri LE. Peptide pheromone-dependent regulation of antimicrobial peptide production in Gram-positive bacteria: a case of multicellular behavior. Peptides. 2001;22(10):1579–1596. doi: 10.1016/S0196-9781(01)00493-4.

9. Erez Z, Steinberger-Levy I, Shamir M, Doron S, Stokar-Avihail A, Peleg Y, et al. Communication between viruses guides lysis–lysogeny decisions. Nature. 2017;541(7638):488–493. doi: 10.1038/nature21049.

10. Stokar-Avihail A, Tal N, Erez Z, Lopatina A, Sorek R. Widespread Utilization of Peptide Communication in Phages Infecting Soil and Pathogenic Bacteria. Cell Host & Microbe. 2019;25(5):746–755.e5. doi: 10.1016/j.chom.2019.03.017.

11. Stewart FM, Levin BR. The population biology of bacterial viruses: why be temperate. Theoretical population biology. 1984;26(1):93–117.

12. Maslov S, Sneppen K. Well-temperate phage: optimal bet-hedging against local environmental collapses. Scientific Reports. 2015;5:10523. doi: 10.1038/srep10523.

13. Berngruber TW, Froissart R, Choisy M, Gandon S. Evolution of Virulence in Emerging Epidemics. PLOS Pathogens. 2013;9(3):e1003209. doi: 10.1371/journal.ppat.1003209.

14. Gandon S. Why Be Temperate: Lessons from Bacteriophage lambda. Trends in Microbiology. 2016;24(5):356–365. doi: 10.1016/j.tim.2016.02.008.

15. Mittler JE. Evolution of the Genetic Switch in Temperate Bacteriophage. I. Basic Theory. Journal of Theoretical Biology. 1996;179(2):161–172. doi: 10.1006/jtbi.1996.0056.

16. Wahl LM, Betti MI, Dick DW, Pattenden T, Puccini AJ. Evolutionary stability of the lysis-lysogeny decision: why be virulent? Evolution. 2018;0(0). doi: 10.1111/evo.13648.

17. Abedon ST. Commentary: Communication between Viruses Guides Lysis–Lysogeny Decisions. Frontiers in Microbiology. 2017;8. doi: 10.3389/fmicb.2017.00983.

18. Abedon ST. Look Who Is Talking: T-Even Phage Lysis Inhibition, the Granddaddy of Virus-Virus Intercellular Communication Research. Viruses. 2019;11(10):951. doi: 10.3390/v11100951.

19. Hershey AD. Mutation of Bacteriophage with Respect to Type of Plaque. Genetics. 1946;31(6):620–640.

20. Doermann AH. Lysis and Lysis Inhibition with Escherichia coli Bacteriophage. Journal of Bacteriology. 1948;55(2):257–276.

21. Kourilsky P. Lysogenization by bacteriophage lambda. Molecular and General Genetics MGG. 1973;122(2):183–195. doi: 10.1007/BF00435190.

22. Abedon ST. Selection for bacteriophage latent period length by bacterial density: A theoretical examination. Microbial Ecology. 1989;18(2):79–88. doi: 10.1007/BF02030117.

23. Abedon ST. Selection for lysis inhibition in bacteriophage. Journal of Theoretical Biology. 1990;146(4):501–511. doi: 10.1016/S0022-5193(05)80375-3.

24. Sinha V, Goyal A, Svenningsen SL, Semsey S, Krishna S. In silico Evolution of Lysis-Lysogeny Strategies Reproduces Observed Lysogeny Propensities in Temperate Bacteriophages. Frontiers in Microbiology. 2017;8. doi: 10.3389/fmicb.2017.01386.

25. Hutchison CA, Sinsheimer RL. Requirement of Protein Synthesis for Bacteriophage phiX174 Superinfection Exclusion. Journal of Virology. 1971;8(1):121–124.

26. Susskind MM, Botstein D, Wright A. Superinfection exclusion by P22 prophage in lysogens of Salmonella typhimurium: III. Failure of superinfecting phage DNA to enter sieA+ lysogens. Virology. 1974;62(2):350–366. doi: 10.1016/0042-6822(74)90398-5.

27. McAllister WT, Barrett CL. Superinfection exclusion by bacteriophage T7. Journal of Virology. 1977;24(2):709–711.

28. Kliem M, Dreiseikelmann B. The superimmunity gene sim of bacteriophage P1 causes superinfection exclusion. Virology. 1989;171(2):350–355. doi: 10.1016/0042-6822(89)90602-8.

29. Bondy-Denomy J, Qian J, Westra ER, Buckling A, Guttman DS, Davidson AR, et al. Prophages mediate defense against phage infection through diverse mechanisms. The ISME Journal. 2016;10(12):2854–2866. doi: 10.1038/ismej.2016.79.

30. Little JW, Shepley DP, Wert DW. Robustness of a gene regulatory circuit. The EMBO Journal. 1999;18(15):4299–4307. doi: 10.1093/emboj/18.15.4299.

31. Paepe MD, Taddei F. Viruses’ Life History: Towards a Mechanistic Basis of a Trade-Off between Survival and Reproduction among Phages. PLOS Biology. 2006;4(7):e193. doi: 10.1371/journal.pbio.0040193.

32. Wang IN. Lysis Timing and Bacteriophage Fitness. Genetics. 2006;172(1):17–26. doi: 10.1534/genetics.105.045922.

33. Shao Y, Wang IN. Bacteriophage Adsorption Rate and Optimal Lysis Time. Genetics. 2008;180(1):471–482. doi: 10.1534/genetics.108.090100.

34. Zong C, So Lh, Sepúlveda LA, Skinner SO, Golding I. Lysogen stability is determined by the frequency of activity bursts from the fate-determining gene. Molecular Systems Biology. 2010;6(1):440. doi: 10.1038/msb.2010.96.

35. Cortes MG, Krog J, Balázsi G. Optimality of the spontaneous prophage induction rate. Journal of Theoretical Biology. 2019;483:110005. doi: 10.1016/j.jtbi.2019.110005.

36. Bossi L, Fuentes JA, Mora G, Figueroa-Bossi N. Prophage Contribution to Bacterial Population Dynamics. Journal of Bacteriology. 2003;185(21):6467–6471. doi: 10.1128/JB.185.21.6467-6471.2003.

37. Gama JA, Reis AM, Domingues I, Mendes-Soares H, Matos AM, Dionisio F. Temperate Bacterial Viruses as Double-Edged Swords in Bacterial Warfare. PLOS ONE. 2013;8(3):e59043. doi: 10.1371/journal.pone.0059043.

38. Bull JJ, Cunningham CW, Molineux IJ, Badgett MR, Hillis DM. Experimental Molecular Evolution of Bacteriophage T7. Evolution. 1993;47(4):993–1007. doi: 10.1111/j.1558-5646.1993.tb02130.x.

39. Bull JJ, Badgett MR, Springman R, Molineux IJ. Genome Properties and the Limits of Adaptation in Bacteriophages. Evolution. 2004;58(4):692–701. doi: 10.1111/j.0014-3820.2004.tb00402.x.

40. Bollback JP, Huelsenbeck JP. Clonal Interference Is Alleviated by High Mutation Rates in Large Populations. Molecular Biology and Evolution. 2007;24(6):1397–1406. doi: 10.1093/molbev/msm056.

41. Betts A, Vasse M, Kaltz O, Hochberg ME. Back to the future: evolving bacteriophages to increase their effectiveness against the pathogen Pseudomonas aeruginosa PAO1. Evolutionary Applications. 2013;6(7):1054–1063. doi: 10.1111/eva.12085.

42. Broniewski JM, Meaden S, Paterson S, Buckling A, Westra ER. The effect of phage genetic diversity on bacterial resistance evolution. The ISME Journal. 2020;14(3):828–836. doi: 10.1038/s41396-019-0577-7.

43. Kerr B, Neuhauser C, Bohannan BJM, Dean AM. Local migration promotes competitive restraint in a host–pathogen ‘tragedy of the commons’. Nature. 2006;442(7098):75–78. doi: 10.1038/nature04864.

44. Heilmann S, Sneppen K, Krishna S. Sustainability of Virulence in a Phage-Bacterial Ecosystem. Journal of Virology. 2010;84(6):3016–3022. doi: 10.1128/JVI.02326-09.

45. Berngruber TW, Lion S, Gandon S. Spatial Structure, Transmission Modes and the Evolution of Viral Exploitation Strategies. PLOS Pathogens. 2015;11(4):e1004810. doi: 10.1371/journal.ppat.1004810.

46. Zinder ND. Lysogenization and superinfection immunity in Salmonella. Virology. 1958;5(2):291–326. doi: 10.1016/0042-6822(58)90025-4.

47. Bailone A, Devoret R. Isolation of ultravirulent mutants of phage Lambda. Virology. 1978;84(2):547–550. doi: 10.1016/0042-6822(78)90273-8.

48. Scott JR, West BW, Laping JL. Superinfection immunity and prophage repression in phage P1 IV. The c1 repressor bypass function and the role of c4 repressor in immunity. Virology. 1978;85(2):587–600. doi: 10.1016/0042-6822(78)90463-4.

49. Hynes AP, Moineau S. Phagebook: The Social Network. Molecular Cell. 2017;65(6):963–964. doi: 10.1016/j.molcel.2017.02.028.

50. Platt TG, Fuqua C. What’s in a name? The semantics of quorum sensing. Trends in microbiology. 2010;18(9):383–387. doi: 10.1016/j.tim.2010.05.003.

51. Popat R, Cornforth DM, McNally L, Brown SP. Collective sensing and collective responses in quorum-sensing bacteria. Journal of The Royal Society Interface. 2015;12(103):20140882. doi: 10.1098/rsif.2014.0882.

52. Silpe JE, Bassler BL. A Host-Produced Quorum-Sensing Autoinducer Controls a Phage Lysis-Lysogeny Decision. Cell. 2018; doi: 10.1016/j.cell.2018.10.059.

53. Ghosh D, Roy K, Williamson KE, Srinivasiah S, Wommack KE, Radosevich M. Acyl-Homoserine Lactones Can Induce Virus Production in Lysogenic Bacteria: an Alternative Paradigm for Prophage Induction. Applied and Environmental Microbiology. 2009;75(22):7142–7152. doi: 10.1128/AEM.00950-09.

54. Rossmann FS, Racek T, Wobser D, Puchalka J, Rabener EM, Reiger M, et al. Phage-mediated Dispersal of Biofilm and Distribution of Bacterial Virulence Genes Is Induced by Quorum Sensing. PLOS Pathogens. 2015;11(2):e1004653. doi: 10.1371/journal.ppat.1004653.

55. Laganenka L, Sander T, Lagonenko A, Chen Y, Link H, Sourjik V. Quorum Sensing and Metabolic State of the Host Control Lysogeny-Lysis Switch of Bacteriophage T1. mBio. 2019;10(5):e01884–19. doi: 10.1128/mBio.01884-19.

